# A framework for peptide identification on commercial nanopore sequencing platforms

**DOI:** 10.64898/2026.05.19.726067

**Authors:** Denis Beslic, Martin Kucklick, Eric Graap, Somayyeh Sedaghatjoo, Bernhard Y. Renard, Stephan Fuchs, Susanne Engelmann, Nils Körber

## Abstract

Direct single-molecule peptide analysis could in principle enable rapid and sensitive identification of pathogen-derived or disease-associated biomarkers without reliance on mass spectrometry. However, existing nanopore peptide sensing methods are typically constrained by limited throughput and lack of accessibility beyond specialized setups.

Here, we present an integrated experimental-computational framework for DNA-linked peptide translocation on a commercially available, high-throughput nanopore sequencing platform, the MinION. Synthetic peptides were covalently bound to oligonucleotides at both termini. The resulting peptide-DNA constructs were then translocated through the CsgG-CsgF pores using a DNA motor protein. Current traces were segmented using the known DNA sequences to extract peptide-associated signal regions. From these segments, we extracted signal features and trained feature-based and deep-learning classifiers to distinguish peptides, balancing interpretability and classification performance.

We establish a framework for peptide identification using standard nanopore sequencing hardware. Across a diverse panel of synthetic peptides, our approach resolves single-amino-acid substitutions, maintains performance across independent sequencing runs, and correctly identifies peptides in blind mixtures. Interpretable model analyses connect classifier decisions and common errors to specific signal motifs. By combining commercially available instrumentation with a reproducible experimental and computational workflow, this framework lowers the barrier to nanopore-based proteomics and enables broader adoption across laboratories. It provides a foundation for future developments in amino acid modification detection and sequence analysis.

## Introduction

Proteins are the key players of life being involved in the vast majority of cellular interactions and reactions. Identifying proteins and changes in their abundance offers valuable insights into essential cellular processes that cannot be fully captured by genome and transcriptome analyses alone. Therefore, analyses based on protein sequencing methods are of central importance for biological and medical research - from basic studies of cellular regulation to the discovery of biomarkers, drug development, and personalized medicine. Advances in mass-spectrometry (MS) based proteomics over the past decades have transformed proteome-scale analysis, enabling the high-throughput identification of thousands of proteins from complex samples ^1^. However, MS-based workflows are typically resource-intensive, involve extensive sample preparation, and can face practical challenges when analyzing highly complex samples or rare modification states ^2,3^. Together, these limitations have driven interest in alternative single-molecule methods that could simplify sample handling and broaden the accessible range of applications in the long term ^4–6^.

Nanopore technology provides an alternative approach to analysing proteins and peptides, offering a route to address these challenges. In a nanopore experiment, an analyte is passed through a nanoscale pore in an insulating membrane under an applied electric potential. The resulting changes in ion current are recorded to derive the local molecular structure. This principle has been translated into real-time, long-read, and modifications-aware sequencing for DNA and RNA on commercially available platforms, with advantages including label-free detection, minimal sample preparation, adaptive sampling, rapid throughput, and single-molecule resolution ^7^. Transferring nanopore methods to proteins and peptides could offer similar benefits for proteomics as it enables the direct, single-molecule reading of amino acid composition and chemical modifications.

While nanopore sequencing is well-established with nucleic acids, applying it to proteins is considerably more challenging. Proteins and peptides are chemically and structurally more heterogeneous: they are composed of 20 canonical amino acids, which give rise to a wide variety of side chains that differ in size, polarity, and charge. Moreover, their higher-order folding can prevent uniform translocation through the nanopore, making controlled unfolding and stepwise movement through the pore a major hurdle ^8,9^. The key difference between peptides and nucleic acids is that peptides lack a uniform, negatively charged backbone that enables helicase-driven, single-residue stepping during DNA sequencing ^3^. Achieving controlled translocation of proteins therefore requires additional strategies, for example through motor enzymes ^10–13^, charged tags ^14–17^, or electro-osmotic flow-based transport ^14,18,19^.

Despite these obstacles, proof-of-concept studies have shown that proteins, peptide fragments and short tagged peptides can generate reproducible ionic-current signatures, indicating that informative structural features are encoded in the signal ^20,21^. These studies have typically relied on custom nanopore setups, motivating efforts to translate these findings to high-throughput commercial platforms through distinct translocation control strategies. Motone and colleagues describe an approach that uses biological unfoldases, such as the ClpX-mediated system, which threads protein strands through CsgG pores of R9.4 MinION flow cells ^13^. While powerful, this strategy depends on ClpX-mediated unfolding and engineered adaptor sequences, which may limit its applicability to native protein structures ^22,23^. A separate line of work uses DNA-peptide conjugates, in which a DNA helicase mechanically pulls the peptide through the pore ^24–26^. However, such DNA-guided methods have been restricted to low-throughput, custom nanopore setups. Recent work by Wang and colleagues demonstrates that high-throughput peptide discrimination is achievable using a custom-engineered nanopore device, but such systems are yet not broadly accessible to most laboratories^27^. Hence, a key outstanding challenge is the adaptation of controlled peptide sensing strategies to widely available nanopore sequencing platforms.

Here, we present a framework for DNA-mediated peptide identification on commercially available sensor arrays. Building on prior research on DNA-peptide constructs for helicase-driven translocation ^11,28,29^, we have adapted the sample preparation and linkage chemistry to enable a robust application on the Oxford Nanopore Technologies (ONT) MinION platform, a widely deployed sequencing system used across thousands of laboratories. Using R10.4.1 flowcells we extracted signal features from the peptide-derived current traces and trained a Light Gradient Boosting Machine (LightGBM) ^30^ for interpretable classification. In parallel, we applied the deep learning-based time-series classifier InceptionTime ^31^ for an improved classification accuracy. To cover a broad range of sequence properties, we designed a panel of 18 synthetic peptides that vary systematically in length, net charge, and hydrophobicity. Across 30 sequencing runs, we collected more than 700,000 individual events, providing a rich dataset for model development and evaluation. We demonstrate that single-amino-acid substitutions yield reproducible and distinguishable signal patterns, and validate classification robustness using blinded mixtures of peptide samples. Finally, we applied model interpretability analyses to link classification decisions to specific signal features and temporal patterns, providing insight into the signal determinants of peptide discrimination and guiding future methodological improvements.

## Results

### Translocation of peptides through CsgG-CsgF nanopores

We adapted a DNA-mediated translocation approach to enable controlled movement of peptide-oligonucleotide constructs (POCs) through the CsgG-CsgF pore on the ONT MinION device. While previous studies demonstrated DNA-assisted peptide translocation through the MspA pore in the *trans*-to-*cis* direction ^11,28,29^, such an approach is not directly compatible with ONT’s high-throughput devices, which restrict access to the trans chamber. Therefore, we designed POCs for *cis*-to-*trans* threading through R10.4.1 flow cells (Figure 1A-B). Key modifications included repositioning the peptide linker to the DNA template’s 3′ end to enforce 5′-entry, and replacing poly(T) threading sequences with heteropolymeric segments terminating in poly(A) (Supplementary Table 1).

**Figure 1.**
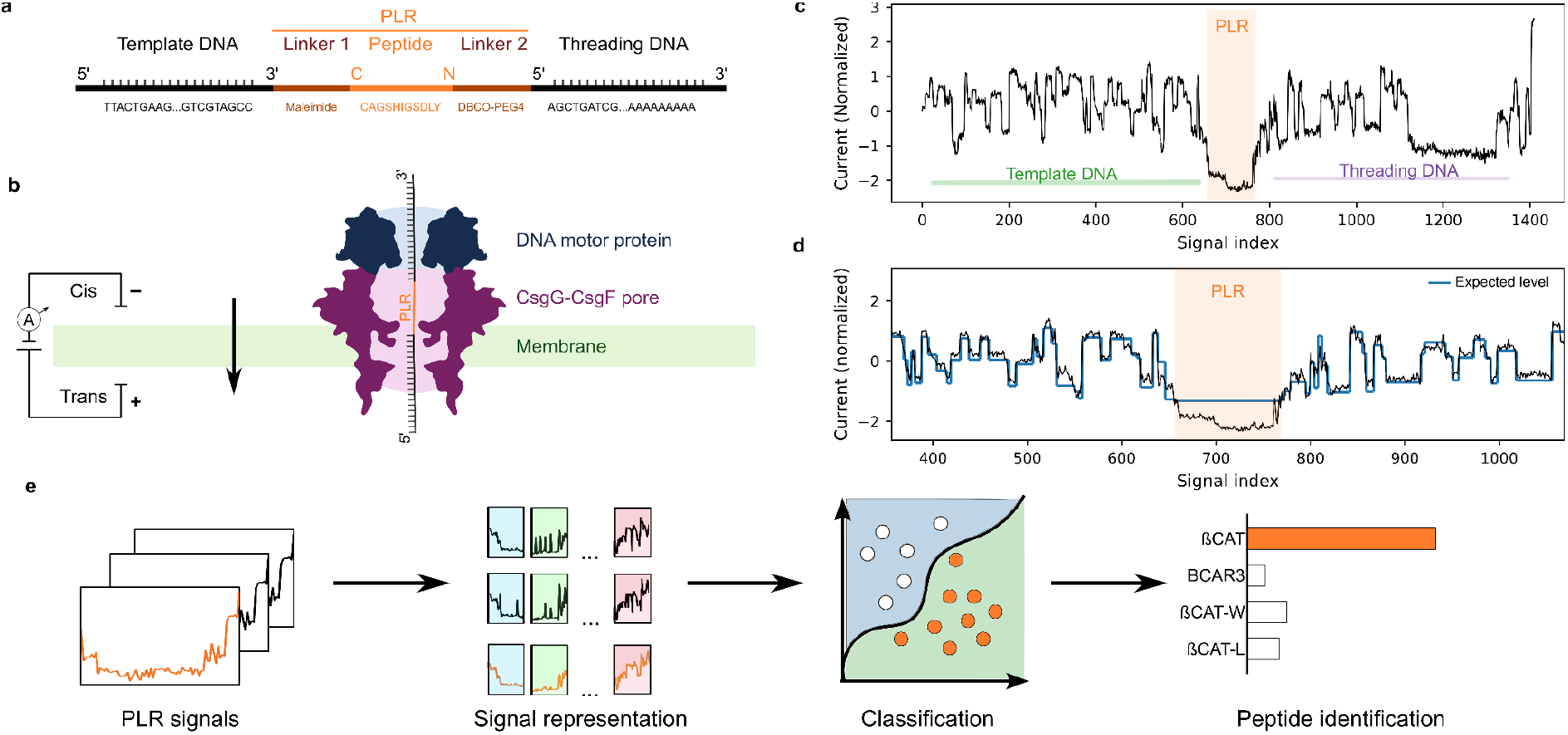
Nanopore translocation of a DNA-peptide-DNA construct through a MinION CsgG-CsgF pore. **(A)** Schematic presentation of the peptide-oligonucleotide construct (POC) comprising the template DNA, a maleimide-thiol linker, the peptide segment, a DBCO-PEG4 linker, and the threading DNA. The peptide-linker region (PLR) encompasses the peptide and its two linkers and is labeled as a single structural unit. **(B)** Illustration of cis-to-trans translocation of the POC through the CsgG-CsgF nanopore, driven by a DNA motor protein. **(C)** An example ionic current trace for a POC containing the ßCAT peptide. Segments that map to the aligned template DNA and threading DNA are indicated by green and violet horizontal bars, respectively, as determined from the basecaller move table via signal-to-reference alignment. The PLR is highlighted in orange. **(D)** Magnified view of the PLR region. Raw current (black) and the expected level inferred from the basecaller move table (blue) are shown. The PLR (orange) is identified by a characteristic drop in current amplitude together with a deviation from the expected DNA-like level pattern. **(E)** Conceptual illustration of how characteristic signal patterns from PLRs are used to assign peptide identity.

As an initial target peptide, we selected the clinically relevant immunopeptide ßCAT. Its physicochemical properties have been shown to support efficient translocation in a previous DNA-linked peptide threading setup ^28^, making ßCAT a practical starting point for adapting the workflow to the CsgG pore on the ONT MinION platform. Following the method described by Nova and colleagues ^28^, we generated double-conjugated peptide-oligonucleotide constructs (POCs), consisting of a template DNA, a centrally linked peptide, and a threading DNA (Figure 1A). POCs were synthesized using an adapted dual-conjugation strategy based on orthogonal bio-conjugation chemistries and subsequently subjected to ONT ligation-based library preparation (Methods). Sequencing adapters required for motor protein loading were ligated to the 5′ end of the template DNA strand. This enables a directed translocation in which the template DNA strand enters the pore first, followed by the attached peptide and finally the threading DNA (Figure 1B). Both double-stranded and single-stranded template configurations were evaluated (Supplementary Table 11), with single-stranded constructs yielding substantially higher numbers of detectable peptide-linker reads and therefore being used in subsequent experiments.

Nanopore current recordings of the ßCAT constructs exhibited low-amplitude current regions occurring between the two DNA segments (Figure 1C). While DNA typically exhibited current levels between 50-165 pA range, these intervals appeared as deeper current reductions (10-50 pA). To investigate the origin of these events, we aligned the raw signals to the reference DNA sequence using basecalling-assisted signal-to-reference mapping (see Methods). Across hundreds of reads, the template DNA alignment terminated immediately before the low-current interval, whereas threading DNA alignment resumed thereafter, matching the expected position of the peptide-linker region (PLR). The predicted DNA-level currents differed substantially from the measured signal within that region (Figure 1D). Based on this consistent alignment pattern and signal deviation, we interpreted the low-amplitude segments as signatures of PLR arising when the conjugated peptide interacts with the pore constriction.

Although the alignment of template and threading DNA around the PLR was observed in a subset of reads, the majority of reads exhibited only alignment to the template sequence. In many cases, basecalls following the template segment did not align to the expected threading sequence. Subsequent analysis revealed that a fraction of these reads instead contained reverse-complement segments of the template within the same read, reflecting a recurrent chimeric configuration independent of peptide identity (Supplementary Note 01). Importantly, the low-current region preceding these segments remained highly consistent in amplitude, duration, and waveform compared to PLRs observed in aligned template-threading reads (Supplementary Fig. 21-23)

Control measurements using only the template DNA produced mostly canonical DNA translocation traces under identical conditions (Supplementary Fig. 3). Only a small number (<50 events) were flagged by the segmentation algorithm (Supplementary Fig. 4). These events were isolated, highly variable, and lacked the reproducible waveform and positional consistency that define peptide-derived PLRs (Supplementary Fig. 1). Thus, while occasional non-specific fluctuations can occur in DNA-only runs, the structured and extended PLR signature was observed exclusively when a peptide was present.

Measurements of the ßCAT POC were reproduced across three independent R10.4.1 flowcells. In all cases, distinct low-amplitude regions downstream of the aligned template DNA were consistently detected (Supplementary Fig. 2), exhibiting highly similar amplitude profiles across replicates.

Together, these results demonstrate that the detected low-amplitude intervals are reproducibly associated with the peptide-linker region passing the pore. Although basecalling accuracy decreases around the PLR due to deviation from DNA-like current profile, integrating coarse basecall information with subsequent signal-level refinement enables consistent localization of the peptide region across reads. These extracted PLR segments form the basis for consensus motif inspection and subsequent supervised classification (Figure 1E).

### Signal Comparison of single-amino-acid variants

Assuming that these detected low-amplitude regions correspond to the peptide, single-amino-acid substitutions within the ßCAT peptide should lead to measurable differences in ionic current due to the different sizes and physicochemical properties of the single amino acids. To test this hypothesis, we synthesized three βCAT variants that differed by one amino acid in the center of the original peptide. The glycine residue at position 5 in the native βCAT was replaced with aspartic acid (βCAT-D), leucine (βCAT-L), or tryptophan (βCAT-W), representing substitutions that vary in size, charge, and hydrophobicity. Each variant was conjugated to the same oligonucleotide sequences and translocated through the nanopore under identical experimental conditions.

All three variants generated ionic current traces resembling the native ßCAT construct: A canonical DNA signal range followed by a low-amplitude interval at the expected position downstream of the aligned template DNA (Supplementary Figure 5). For each variant, a subset of 2,000 PLR signals were aligned using pairwise dynamic time warping (DTW), and a representative median signal was chosen based on the smallest DTW distance to all other signals (Figure 2A). The depth and duration of the current reduction varied with the substituted amino acid residue. The bulky tryptophan variant ßCAT-W generated the most pronounced reduction in current, while ßCAT-D and ßCAT-L exhibited subtler shifts. Comparison of signal point distributions across the 2,000 aligned PLRs revealed generally lower median current levels for ßCAT-W and ßCAT-L compared to ßCAT and ßCAT-D (Supplementary Figures 7).

Because simple comparisons of median and standard deviation values did not fully capture the differences between variants, we computed a comprehensive set of 50 manually selected features covering statistical, shape-based, and temporal properties, including slope and derivative metrics, peak characteristics, energy and trend measures, and catch22 time-series features (Figure 2B, Supplementary Note 4). Feature-level comparisons between ßCAT and ßCAT-W revealed systematic differences primarily in slope- and derivative-related metrics, reflecting a faster current decline and recovery for ßCAT-W. The aspartate substitution (ßCAT-D) caused only minor deviations from native ßCAT, while the leucine variant (ßCAT-L) showed intermediate behavior between ßCAT and ßCAT-W. At the same time, the median signals and feature distribution of the ßCAT repeated measurements confirmed the reproducibility across different flow cells. Minor visual variations between medoid traces likely stem from subsampling effects and DTW sensitivity to outliers, but the underlying signal statistics remained highly stable across experiments.

Dimensionality-reduced projections (PCA, t-SNE, UMAP) of the 40 most discriminative features supported these observations: ßCAT and ßCAT-D formed overlapping local clusters, whereas ßCAT-L and ßCAT-W occupied more distinct regions of feature space (Figure 2C). Although clear class boundaries did not emerge, these structured local patterns demonstrate that variant-specific signal characteristics could be distinguished and exploited for classification. These results demonstrate that the MinION platform can resolve sequence-dependent current signatures arising from single-amino acid substitutions, providing a foundation for discrimination of peptide identity.

**Figure 2.**
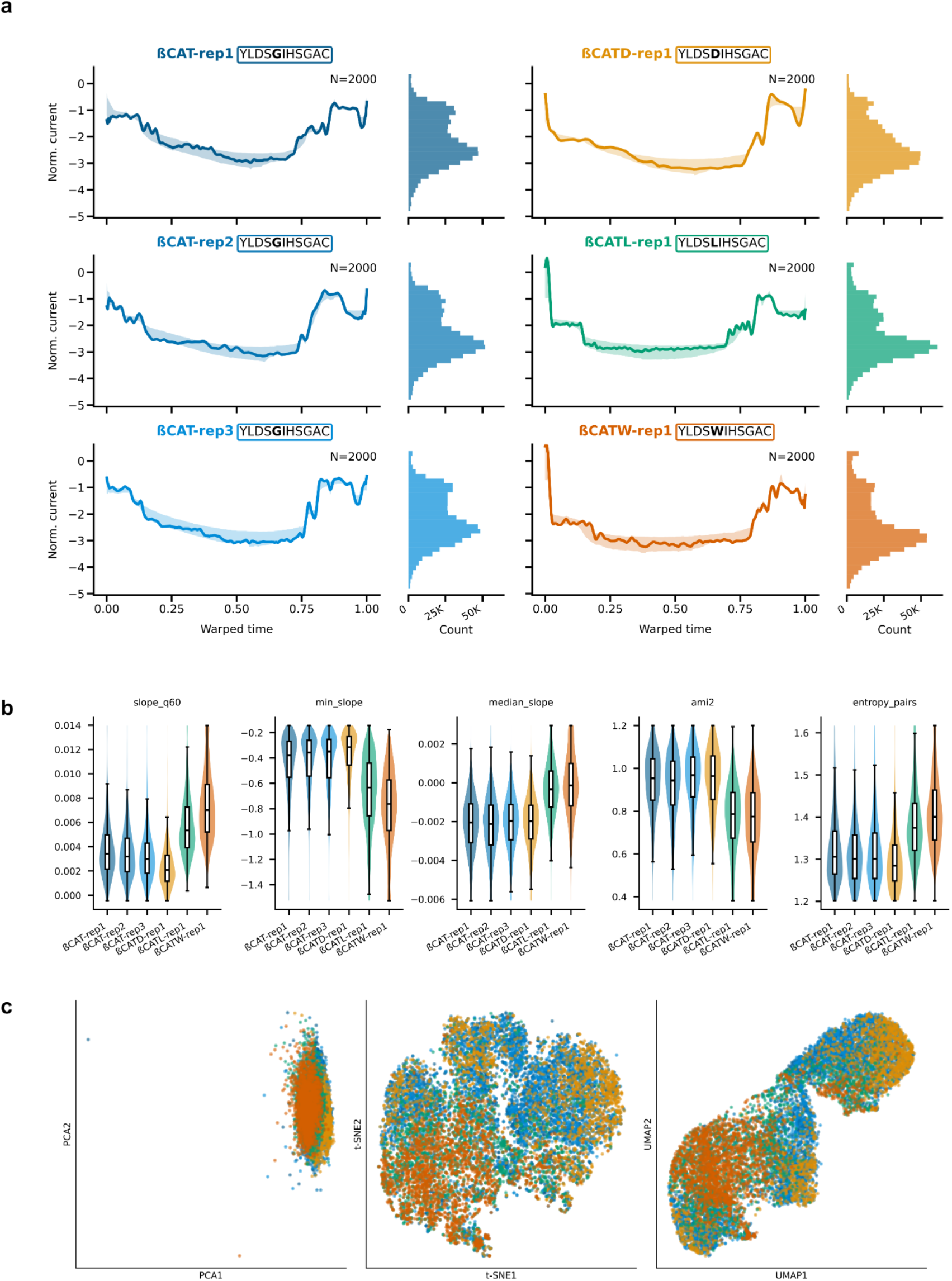
Resolution of single-amino-acid substitutions in ßCAT peptides. **(A)** Representative nanopore current signals for βCAT (blue) and its single-amino-acid variants βCAT-D (yellow), βCAT-L (green), and βCAT-W (orange). The dark solid line shows the DTW median signal, the current trace with the lowest DTW distance to all other signals in the class. Shaded regions denote the interquartile range of all other signals DTW-aligned to the median signal. Adjacent histograms show the distribution of signal values across all traces. **(B)** Box plots showing the distribution of the five most discriminative signal features across variants, highlighting systematic differences among βCAT variants. Values were clipped at 1st-99th percentiles for visualization. **(C)** Two-dimensional projections (PCA, t-SNE, UMAP) based on 30 extracted signal features, illustrating class separability among single-amino-acid variants.

### Machine learning-based classification of peptide-specific current signals

To determine whether nanopore current traces contain sufficient information for peptide identification, we developed a machine learning framework, including PLR extraction, feature extraction, model training, and evaluation. From each read we extracted the PLR using the alignment-anchored changepoint procedure, followed by normalization, low-pass filtering, and resampling to a fixed length (see Methods). Classification performance was primarily assessed using the deep learning-based time-series classifier InceptionTime ^31^, which consistently achieved the highest accuracy across datasets. In parallel, we performed feature-based classification using a gradient-boosted decision-tree model trained on a 50-dimensional feature set to enable interpretability of signal characteristics. For comparison, we also evaluated alternative convolution-based and end-to-end deep learning approaches (Supplementary Table 5), which are typically used for time-series classification ^32^.

As an initial step, classification was performed on four ßCAT variants (ßCAT, ßCAT-D, ßCAT-L, ßCAT-W), which differ only in their amino acid sequence at a single central position. We trained on 90% of reads pooled from multiple sequencing runs and validated on the remaining 10%, while fully independent runs served as the final test set (Figure 3C; Methods). In this four-class task, InceptionTime achieved an accuracy of 92.3% and 92.1% on the validation and held-out test sets, respectively, with macro-averaged F1 scores of 88.9% and 91.8% (Fig. 3A-B).

**Figure 3.**
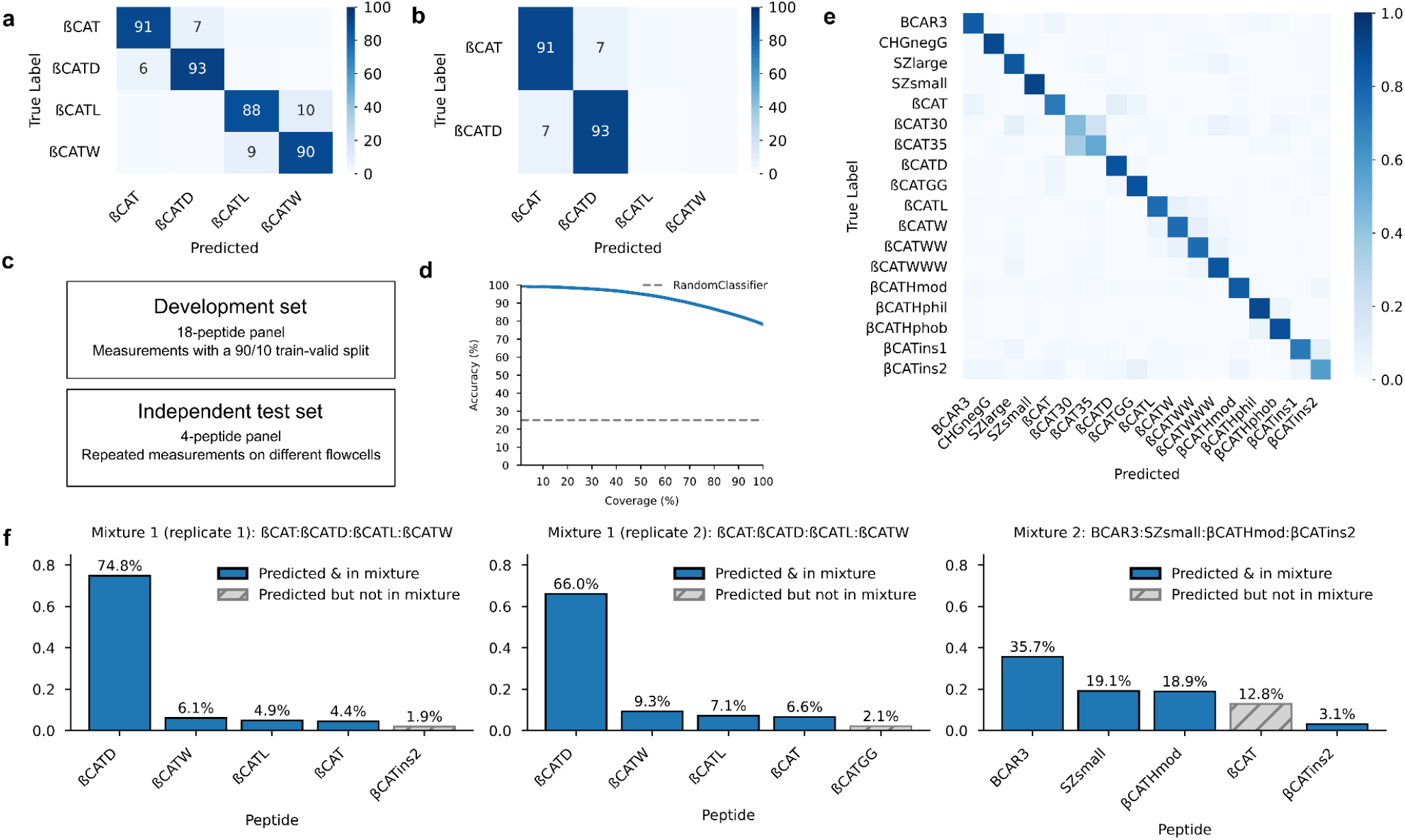
Machine learning-based peptide classification from nanopore current traces using InceptionTime. **(A)** Confusion matrix for the validation dataset of the four-peptide benchmark. **(B)** Confusion matrix for the independent test dataset of the four-peptide benchmark. For clarity, only cells accounting for >10% of all predictions are shown. **(C)** Overview of dataset partitioning. The development dataset comprised measurements from 18 peptides used for model training and validation, while the independent test dataset included measurements from four peptides. **(D)** Recall-coverage curve for predictions on the full test dataset. The light grey dotted line indicates the expected performance of a random classifier. **(E)** Confusion matrix for the complete validation dataset across all peptide classes. **(F)** Predicted composition of the blind peptide mixtures. For each mixture and replicate, the classifier’s highest-confidence predictions are shown as bar plots. Peptides present in the ground-truth mixture are highlighted in blue, whereas off-target predictions are shown in grey with hatching.

Next, we extended the task by including βCAT-GG (two glycines), βCAT-WW (two tryptophans), and βCAT-WWW (three tryptophans). This seven-peptide panel expansion posed a markedly more difficult classification problem, as all three W-containing peptides exhibited similarly shaped deep current profiles (Supplementary Figure 11). While amplitude- and slope-based feature distributions overlapped considerably, InceptionTime achieved strong performance, achieving accuracies of 90.0% on the validation set and 87.5% on the test set, with corresponding macro F1 scores of 84.9% and 91.1%. Misclassifications were largely confined to the tryptophan-rich group (Supplementary Fig. 13).

We next extended the analysis to a diverse panel of 18 peptides spanning 11 to 35 residues in length, with net charges ranging from −7 to +2 and broad hydrophobicities (Supplementary Table 2-3). Two highly positively charged peptides (CHG-posD, CHG-posL) were excluded due to low PLR yields (< 200) and excessive noise in the signals. The efficiency of PLR detection varied between constructs, with some (e.g. ßCAT, BCAR3) yielding over 10,000 segments and others (e.g. ßCAT-35) generating less than 1,000. This imbalance correlated with reduced template DNA alignments counts (Supplementary Table 4), suggesting a lower number of successful translocation events rather than segmentation errors. Nevertheless, all 18 peptides exhibited low-current events downstream of the template DNA alignment that consistently deviated from DNA move-table predictions (Supplementary Fig. 5-6).

On the full 18-peptide panel, InceptionTime achieved accuracies of 83.1% in the validation set and 78.2% on the independent test set, with macro F1 scores of 63.7% and 74.0%, respectively (Figure 3D-E). Applying a confidence threshold enabled further precision-coverage tradeoffs: restricting predictions to examples with ≥75% classifier confidence increased accuracy on the validation set to 96.1% but reduced coverage to 63.9% of PLRs. Consistent with results from the 7-peptide panel, most misclassifications occurred among the βCAT-W/WW/WWW variants due to their similar low-amplitude shapes. Reduced accuracies for long βCAT constructs (βCAT-30/35) likely stem from elevated noise (Supplementary Fig. 6) and a smaller number of training examples (Supplementary Table 4). The lower performance of insertion variants (βCAT-ins1/2) may also reflect poorer purification quality (Supplementary Table 2), which may have introduced additional signal variability.

To evaluate our approach in an application context, we applied the trained classifier to unknown peptide mixtures. Mixture 1 contained βCAT, βCAT-D, ßCAT-W, and ßCAT-L. Mixture 2 contained BCAR3, SZ-small, βCAT-Hmod, and ßCAT-ins2. Each mixture was prepared at a ratio of 1:1:1:1. However, the effective representation of each peptide at the signal level is not expected to strictly reflect the input stoichiometry. Differences in POC formation efficiency, adapter ligation, and peptide-specific physicochemical properties influencing translocation through the pore can introduce systematic biases, leading to unequal representation in the recorded reads. Hence, the analysis focuses on qualitative identification of mixture components rather than precise quantification. For Mixture 1, the classifier identified the correct peptides as the most frequently predicted classes across the two replicates (Figure 3G). For Mixture 2, three of the four most frequently predicted classes matched the ground truth, with the single misclassified peptide ßCAT-ins2, a peptide that was already partially confounded with βCAT in the validation dataset. While the model does not reproduce exact quantitative mixing ratios, it robustly recovers the dominant constituents in both mixtures, demonstrating correct peptide identification in complex samples.

To investigate which signal characteristics drive model decisions, we performed interpretability analyses on the feature-based LightGBM model, as its explicit feature representation enables direct attribution. While InceptionTime achieved a higher predictive performance, its internal representations are less directly interpretable. We combined SHAP (SHapley Additive exPlanations)-based global and local importance with a sliding window occlusion analysis (Methods). Global SHAP rankings revealed that slope-derived features dominated model decisions, highlighting that the rate and trajectory of current drop and recovery encode important information for peptide classification (Figure 4A). Local SHAP analyses illustrated feature contributions for class-specific predictions: short decreasing stretches and rapid slope changes strongly supported assignments to ßCAT-W and ßCAT-L, whereas smoother, monotonic decreases provide stronger probability for ßCAT and ßCAT-D (Figure 4B). Complementary temporal occlusion analysis confirmed the importance of slope dynamics and showed that signal segments near the PLR start and end are particularly important for classification (Figure 4C). This pattern suggests that there may be segmentation or alignment errors at PLR boundaries. Unaccounted DNA segments could influence both extracted features. We extended this analysis to the tryptophan-containing variants of ßCAT and observed similar patterns of importance, although the overlapping blockade shapes in these constructs reduce discriminability and concentrate misclassifications within the tryptophan-rich subgroup (Supplementary Figure 14).

**Figure 4.**
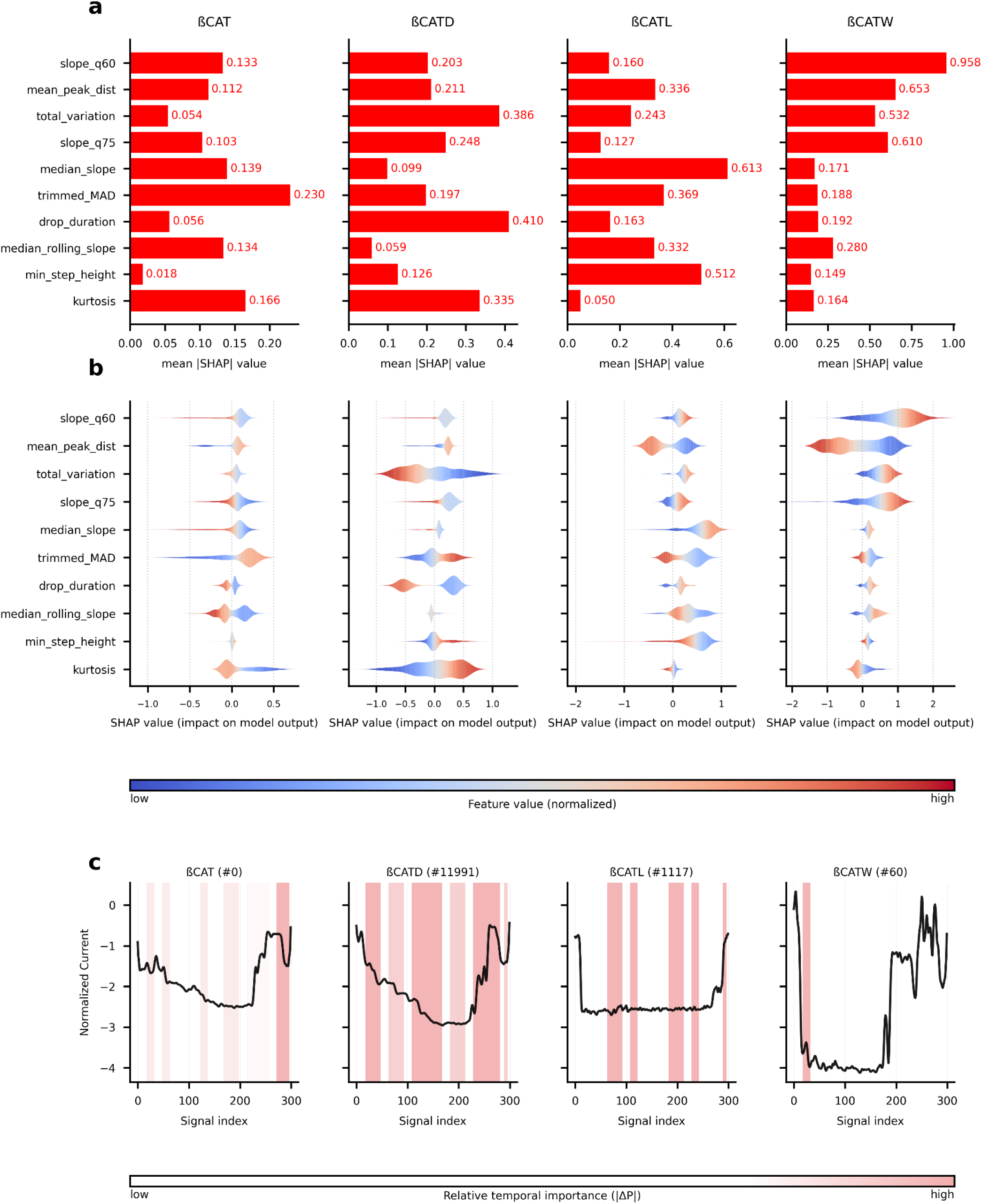
Model interpretability and error analysis on the LightGBM model. Each column corresponds to one peptide (ßCAT, ßCAT-D, ßCAT-L and ßCAT-W), showing peptide-specific global, local and temporal feature importance. **(A)** Global SHAP feature importance showing the mean absolute SHAP value of the ten most influential features for each peptide, based on the global feature ranking across all classes. Higher values indicate stronger overall contribution to the model’s predictions. **(B)** Local SHAP feature importance visualized as beeswarm plots for the same set of features. Each point representModes a single prediction, where color denotes the corresponding normalized feature value (blue for low, red for high). Positive SHAP values indicate a feature increases the predicted probability for a given peptide, while negative values indicate the opposite effect. **(C)** Temporal feature importance for representative examples. Black traces show the raw nanopore current signals, and red-highlighted regions mark segments whose occlusion (replacement with current trace median) causes the largest change in predicted class probability. Color intensity reflects the magnitude of |ΔP| and is normalized separately for each example to represent the relative occlusion effect within that signal.

## Discussion

We introduced a scalable experimental-computational framework for peptide detection and classification on widely available nanopore sequencing platforms. Peptides were linked to oligonucleotides and translocated through CsgG-CsgF pores with a DNA motor protein. This generated PLR current traces, which contain peptide-specific signal patterns. Extracting a comprehensive set of signal features from these PLRs and applying machine-learning classification enables us to resolve single amino-acid substitutions, distinguish synthetic peptides with different sequences, and identify the dominant components of mixed peptide samples. Together, this demonstrates that standard nanopore sequencing devices can be repurposed for peptide analysis without hardware modification.

Compared with previous studies on nanopore-based peptide and protein sensing, our work expands both accessibility and ability to analyze peptides with challenging physicochemical characteristics. Earlier approaches using aerolysin ^33,34^, FraC ^35^, or engineered MspA variants ^36,37^ have demonstrated impressive sensitivity to individual amino acids and post-translational modifications. However, these studies typically rely on controlled, low-throughput single-pore measurements, specialized pore engineering, or custom instrumentation, that limit scalability and accessibility. Recent work extends single-amino-acid resolution to commercial MinION devices using ClpX-mediated threading on R9.4 flow cells ^13^. In contrast, we analyzed DNA-linked peptides moved by the standard DNA motor protein through R10.4 pores. Our approach provides a distinct route to peptide-level readout on unmodified, commercially available flow cells. At the same time, a recent study by Wang and colleagues demonstrates high-resolution peptide discrimination using custom-engineered high-throughput sensors ^27^. While this highlights the potential of optimized, specialized devices, such systems are not broadly accessible. Our approach instead focuses on deployability by leveraging widely available MinION devices, enabling massively parallel measurements on thousands of pores simultaneously and systematic exploration of more challenging peptide groups, including those with varying lengths and sets of near-identical sequences. Despite the increased complexity and reduced signal separability of these constructs, we are able to recover class-specific patterns, detect single-residue substitutions, and identify dominant components in peptide mixtures. Large-scale measurements further reveal limitations that are less apparent in low-throughput approaches, including sequence-dependent differences in translocation efficiency, as well as physicochemical biases associated with charge, length, and hydrophobicity. These biases affect both PLR yield and signal quality. Overall, these findings establish our study as a bridge between specialized proof-of-concept demonstrations and a deployable, high-throughput framework for nanopore-based peptide analysis on standard sequencing platforms.

While the framework is already deployable on standard sequencing hardware, several technical constraints currently restrict its broader applicability. First, our conjugation scheme relies on a C-terminal cysteine for thiol-maleimide coupling, restricting the range of compatible peptides and complicating the analysis of proteolytic digested peptides. Developing ligation chemistries that accommodate diverse termini and side chains will therefore be essential for proteome-scale applications. Second, peptide translocation efficiency and signal quality depends on charge, length, and hydrophobicity. In particular, highly cationic or longer peptides often exhibit inefficient capture or incomplete translocation. These biases were also reflected in the mixture experiments, where specific constructs yielded fewer PLR events despite comparable concentrations. Overcoming these limitations through optimized linker design, voltage control, and buffer composition will be key to achieving uniform detection across diverse peptide classes ^23,38^. The physical dimensions of the nanopore add another layer of complexity. Shorter pores, such as MspA or CsgG, offer higher spatial resolution and could better resolve heterogeneous peptide sequences. However, their limited length constrains the maximum readable peptide length to around 25 amino acids ^12^, which restricts analysis of longer peptides and protein fragments. Our use of CsgG-CsgF pores with a longer dual-constriction structure ^39^, extends the reading length but may reduce resolution. Future improvements in nanopore design and control of translocation speed will be critical to balancing resolution and peptide length, enabling both sequence discrimination and analysis of longer peptides.

Another major source of variability originates from the PLR segmentation. While the changepoint-based algorithm reliably identifies PLRs, slight misalignments at the DNA-peptide border can lead to inconsistent boundary assignment and signal misclassification. Incorporating short, well-defined DNA homopolymers that flank the peptide could provide clearer landmarks for segmentation, enabling improved boundary precision. Similarly, expanding the training dataset to include multiple flow cells and replicate runs will be essential to capture instrument-specific variability and improve model generalizability. Furthermore, our work focuses exclusively on synthetic peptides, and performance on complex biological samples remains to be established.

From a computational perspective, our workflow establishes a first step toward peptide identification using integrated signal processing and machine learning on nanopore sequencing data. Taking this approach further to enable residue-level peptide decoding would require substantially larger and more systematically designed training sets, as well as improved biophysical models of peptide-pore interactions. Furthermore, advances in motor control, pore engineering, and adaptive voltage protocols will be necessary to slow and regularize peptide motion through the pore. This will enhance the signal-to-noise ratio and temporal resolution ^38^.

Another technical consideration is the occurrence of chimeric forward-reverse complement template reads. In a subset of reads, the expected template-threading configuration was replaced by a continuous translocation event comprising a forward template segment followed by its reverse complement. Multiple lines of evidence suggest this configuration is most consistent with library preparation-associated fold-back or internal priming, potentially mediated by secondary structure formation and polymerase-driven strand extension (Supplementary Note 01). Similar artefacts have previously been described in long-read sequencing and enzymatic library preparation workflows ^40,41^. Importantly, the peptide-linker signal precedes the transition and remains unchanged across configurations. Accordingly, classification performance is unaffected, as confirmed by cross-configuration analyses (Supplementary Fig. 17-18). These findings indicate that the chimeric reads represent reproducible library-associated artefacts that do not affect peptide detection or principal conclusions of this study.

Despite these challenges, the demonstrated ability to perform peptide detection and classification on unmodified, commercially available nanopore sequencing devices represents an important step towards accessible and scalable peptide analytics. Continued progress in instrumentation, chemistry, and machine learning, could see nanopore-based peptide assays complementing LC-MS/MS in applications where portability, cost-efficiency, or single-molecule resolution are key priorities ^42^. Benchmarking against established proteomic workflows will be crucial in assessing quantitative accuracy and dynamic range. At the same time, targeted assays and multiplexed DNA barcoding may enable early practical applications ^43^.

More broadly, our results demonstrate that nanopore-based peptide analysis can be realized on existing high-throughput sequencing platforms without requiring specialized hardware modifications. By combining a compatible experimental design with a scalable computational workflow, this approach lowers the barrier to entry for nanopore proteomics and enables its adoption across a wide range of laboratories. In this way, our work provides a foundation for extending nanopore sequencing platforms, originally developed for nucleic acids, to peptide analysis.

## Methods

### Preparation of Peptide-Oligonucleotide Conjugates

Peptides were synthesized by JPT Peptide Technologies and GenScript with a manufacturer-verified purity of 90% (analytical RP-HPLC and LC-MS). Due to challenges in the synthesis of complex or hydrophobic sequences, a subset of peptides was supplied with crude purity (Supplementary Table 1). DNA oligonucleotides (HPLC-purified) were purchased from Biomers (Supplementary Table 2).

Conjugates were assembled using a dual-chemistry strategy adapted from established protocols ^11,28^: Peptides (N-terminal azide, C-terminal cysteine) were mixed with 5′-DBCO-PEG4 modified threading DNA and 3′-maleimide modified template DNA at a molar ratio of 1:2:6 (peptide:threading DNA:template DNA) in a phosphate buffered saline (PBS, pH 7.4). The peptide concentration was adjusted to 7 µM (total volume 30-45 µL). Reactions were incubated for 20 hours under anaerobic conditions and 4 °C to limit side reactions and cysteine oxidation.

Following conjugation, 8µL of the conjugate mixture were desalted using the oligonucleotide cleanup protocol of a Monarch DNA Gel extraction Kit (New England Biolabs, #T1020L). The mixture was further purified using the NEB High Input Poly(A) mRNA Isolation Module (NEB, #E3370). Poly-T beads selectively captured the poly-A-tagged POCs, removing non-bound template DNA, peptides, and other residual components.

### Nanopore Sequencing protocol

All nanopore measurements were performed on the Oxford Nanopore Technologies (ONT) MinION Mk1B device with R10.4.1 flow cells (FLO-MIN114). Flow cells were primed and operated in standard ONT running buffers. The data acquisition was performed with the MinKNOW software (v23.04.6+) at 180 mV applied voltage and 5 kHz sampling rate.

Sequencing libraries were prepared using the Ligation Sequencing Kit V14 (ONT, SQK-LSK114), following the manufacturer’s protocol (genomic-dna-by-ligation-sqk-lsk114-GDE_9161_v114 _revO_29Jun2022-minion). The volume of the POCs was adjusted to 47 µL with nuclease-free water. DNA repair and end-prep were performed using NEBNext FFPE DNA Repair Mix (NEB, #M6630) and the NEBNext Ultra II End Repair/dA-Tailing Module (NEB, #E7546) with a DNA control sample which was included in the kit. Adapter ligation was performed using the NEBNext Quick Ligation Module (NEB, #E6056), and ligated libraries were purified using Short Fragment Buffer. The resulting eluate was used as the final sequencing library.

Libraries were sequenced on a MinION Mk1B device with R10.4.1 flow cells (ONT, FLO-MIN114). Flow cells were primed and loaded with a final POC concentration of ∼750 nM, unless otherwise stated in Supplementary Table 3. Data acquisition and run control were performed using MinKNOW v23.04.6+, and sequencing runs lasted 48-72 hours depending on the experiment.

### Nanopore Data analysis

#### Basecalling and alignment

Raw current signal data (.pod5) were basecalled using the Dorado software (v1.3.1) ^44^ with the “dna_r10.4.1_e8.2_400bps_sup” model. The “--emit-moves” parameter was enabled to retain move-table (mv) tags, ensuring preservation of base-to-signal coordinate mappings alongside basecall sequences and quality scores. Basecalled reads were aligned with minimap2 (v2.26+) ^45^ with adjusted ONT-specific parameters (-ax map-ont -k 9 -w 5 --secondary = no). Alignments were performed against a concatenated reference sequence containing the template DNA (Supplementary Note 2).

Since move-table (mv) tags emitted by the basecaller provide only coarse signal-to-base mappings prone to alignment inaccuracies, we refined these mappings using remora.io utilities (v0.6.0) ^46^. The signal mapping refinement (re-squiggling) step improves move-table segmentation and allows comparisons between different reads (see Supplementary Note 1+3).

#### Segmentation

PLRs were identified per read using an alignment-guided signal segmentation approach. Raw ionic current signals were extracted from POD5 files and aligned to the reference sequence. The reference coordinate corresponding to the end of the template DNA served as an anchor point; if this exact position was not aligned, the nearest preceding aligned reference position was used. A restricted signal window centered on the anchor (±70 bases, converted to signal coordinates using the median query-to-signal stride) was used for segmentation.

Within this window, the DNA baseline current was estimated as the median of the upstream signal (first 25% of the window, up to 1,000 samples). Candidate peptide-linker regions were detected as sustained drops in current relative to this baseline. The baseline-subtracted signal was smoothed using uniform filters (15 and 35 samples), and samples exceeding a drop threshold (≥45% of the minimum pA-drop criterion, minimum 5 pA) were marked as candidate regions.

For each candidate region, the current drop (difference between the DNA baseline and the 25th percentile signal within the region) and the deviation from the expected move-level current were computed. Regions were required to have length ≥60 samples, pA drop ≥75 pA, and move-level deviation ≥15 pA. The highest-scoring region satisfying these criteria and located near the DNA-peptide junction (≤500 signal samples) was selected as the PLR. Boundaries were refined to the nearest query-to-signal indices, and reads with invalid signal or failing quality checks were excluded.

#### Signal processing

To enable direct comparison between reads, extracted PLR signals were normalized with Remora. The signals were smoothed with a 2nd-order low-pass Bessel filter using a cutoff of 0.2 and then resampled to a fixed length of 300 samples. For each sequencing run, all samples were pooled and split into training (90%) and validation (10%) sets using class-stratified random splitting. Independent sequencing runs performed on separate flow cells containing the same peptide standards were used as the test set, ensuring evaluation on unseen runs and confirming reproducibility across flow cells (Supplementary Table 4).

#### Consensus signal generation

Representative consensus traces for each peptide class were obtained using a DTW-based approach ^47^. Up to 2,000 reads per class were uniformly subsampled, and pairwise DTW distances were calculated using a Sakoe-Chiba band (10%) with path normalization. The single trace showing the smallest total distance to all other sampled traces was selected as the class representative. This preserves an actual observed signal (unlike arithmetic averages) and enables direct visual interpretation. All other signals were aligned to the median trace, and the interquartile range of these DTW-aligned signals relative to the median was visualized.

#### Feature extraction and classification

Processed PLR signals were transformed into interpretable features combining statistical, shape-based, and temporal properties. An initial set of 152 time-series features was extracted per signal, including statistical descriptors (mean, median, variance, skewness, kurtosis, MAD), slope- and trend-based measures (e.g., median slope, quantile slopes), signal-drop metrics describing the largest downward transition in the signal, and 22 canonical time-series features from the catch22 library ^48^.

To reduce redundancy and model complexity, 50 informative features were selected from the full feature set based on feature importance ranking on the training data. The resulting feature subset was used for all classification experiments. Details of the feature extraction procedure and the feature selection protocol are provided in Supplementary Note 4, and the selected features are listed in Supplementary Table 13.

Feature-based classification was performed using a LightGBM gradient-boosted tree model (v4.3.0) ^30^ with adaptive class weighting and 250 boosting rounds to achieve stable convergence.

In parallel, deep learning-based classification was performed using InceptionTime as implemented in the aeon library (v1.2.0) ^49^. Model parameters were set to kernel size = 64, depth = 4, n_classifiers = 5, n_filters = 64, n_conv_per_layer = 3, and a learning rate of 1.5 × 10^−3^. Early stopping was applied during training of InceptionTime. Model training was monitored based on the validation loss, with a patience of 50 epochs and a minimum improvement threshold (min_delta) of 0.0. Training proceeded for a maximum of 1,000 epochs and was terminated early if no improvement in validation loss was observed within the specified patience window. The model weights corresponding to the lowest validation loss were restored. Additional benchmarking against alternative time-series classifiers is provided in Supplementary Table 5.

#### Model interpretability

Feature-level interpretability was assessed using SHAP ^50^, applied to the trained LightGBM model on the validation dataset. For global interpretation, we ranked features by their mean absolute SHAP value averaged across all peptide classes and selected the ten most influential features for visualization. These features represent the dominant contributors to the model’s decision space.

Local interpretability was visualized using beeswarm plots constructed for each peptide, restricted to the same globally ranked top-10 features to enable direct cross-class comparison. Points represent individual predictions, where SHAP values indicate direction and magnitude of influence, and point color encodes the normalized raw feature value.

Temporal contributions to classification were evaluated via a sliding-window occlusion analysis. For each misclassified peptide, a moving window (width = 50, step = 20) was replaced by the signal’s median, and the resulting change in predicted probability for the true class (ΔP = P_baseline_ - P_occluded_) quantified local importance. Regions with large positive ΔP values denote segments most critical for classification decisions.

## Data availability

The code used for the nanopore data analysis is available on GitHub at https://github.com/ZKI-PH-ImageAnalysis/nanopore-peptide-identification.

## Author information

## Author contribution

D.B. and M.K. contributed equally to this work. N.K., S.E., and S.F. share senior authorship. D.B, N.K., B.R., S.F., and S.E. conceived the study. D.B. developed the code, performed the nanopore data analysis, and drafted the manuscript. M.K. and E.G. synthesized the DNA-peptide constructs and carried out the nanopore measurements. N.K., B.R., S.F., and S.E. supervised the project. All authors revised the manuscript and approved the final version.

## Author correspondence

Correspondence to Stephan Fuchs, Susanne Engelmann, and Nils Körber.

