## Supplementary Data for "A framework for peptide identification on commercial nanopore sequencing platforms"

### Contents

**Contents** **2**
**Supplementary Notes** **3**
Supplementary Note 01 | Chimeric reverse complement template artefacts 3
Supplementary Note 02 | Signal refinement of nanopore sequencing signals 5
Supplementary Note 03 | PLR segmentation 6
Supplementary Note 04 | Feature extraction overview 7
**Supplementary Tables** **9**
Supplementary Table 1 | Sequences of DNA oligonucleotides and the Peptide-Oligonucleotide Construct. 9
Supplementary Table 2 | Sequences and purification characteristics of the analyzed peptides. 9
Supplementary Table 3 | Physicochemical properties and characteristics of sequenced peptides. 11
Supplementary Table 4 | Summary statistics for MinION nanopore sequencing runs. 12
Supplementary Table 5 | Performance metrics across the full dataset for different models. 15
Supplementary Table 6 | Per-class validation performance metrics for the LightGBM and InceptionTime model. 15
Supplementary Table 7 | Per-class test performance metrics for the LightGBM and InceptionTime model. 16
Supplementary Table 8 | Validation performance of the InceptionTime model with varying numbers of included peptides. 16
Supplementary Table 9 | Sequences of DNA oligonucleotides used for minimap2 alignment 16
Supplementary Table 10 | Alignment statistics using different reference sequences for nanopore sequencing runs. 17
Supplementary Table 11 | Comparison of βCAT single-strand and double-strand variants. 18
Supplementary Table 12 | Top 50 selected features used for classification. 19
**Supplementary Figures** **21**
Supplementary Figure 1 | Segmentation of βCAT signals. 21
Supplementary Figure 2 | Reproducibility of PLR signals. 21
Supplementary Figure 3 | Control experiments with template-only constructs. 22
Supplementary Figure 4 | False-positives from control experiment with template-only constructs. 22
Supplementary Figure 5 | PLR traces from βCAT-D (A), βCAT-L (B), and βCAT-W constructs (C). 23
Supplementary Figure 6 | PLR traces from BCAR3 (A), SZ-large (B), and ßCAT-30 (C). 23
Supplementary Figure 7 | Overlap of signal distributions across βCAT peptide variants. 24
Supplementary Figure 8 | Confusion matrix for peptide mixture experiments. 24
Supplementary Figure 9 | Signal comparison between βCAT and βCAT-W. 25
Supplementary Figure 10 | Signal comparison between repeated measurements of βCAT and βCAT-W. 26
Supplementary Figure 11 | Signal comparison between βCAT, βCAT-W, βCAT-WW, and βCAT-WWW 27
Supplementary Figure 12 | Peptide physicochemical properties and corresponding classification performance metrics. 28
Supplementary Figure 13 | Spearman correlations between peptide physicochemical properties and classification metrics. 28
Supplementary Figure 14 | Model interpretability and error analysis for W-variants of ßCAT. 29
Supplementary Figure 15 | Forward-reverse complement template chimeras identified in βCAT. 30
Supplementary Figure 16 | Forward-reverse complement artefacts in reads aligning exclusively to threading DNA. 30
Supplementary Figure 17 | Comparison of signal and features of template-RCtemplate and template-threading alignments. 31
Supplementary Figure 18 | Classification performance across alignment configurations for ßCAT and ßCAT-WW. 32
Supplementary Figure 19 | Current traces from peptide-free control runs. 32
Supplementary Figure 20 | Secondary structure prediction of the template DNA. 33
Supplementary Figure 21 | Secondary structure prediction of the threading DNA. 34
Supplementary Figure 22 | Secondary structure prediction of the template-threading construct. 35
**References** **36**

### Supplementary Notes

**Supplementary Note 01 | Chimeric reverse complement template artefacts**

**Overview**

During analysis of PLR-containing reads, we observed chimeric basecalls in which the expected forward template sequence was followed by a reverse-complement (RC) segment of the same template within a single continuous read. Because the main manuscript focuses on peptide classification, we provide here a detailed characterization of these artefacts, the control experiments performed, and possible explanations. We demonstrate that this phenomenon is reproducible, occurs independently of peptide presence, and does not affect the central findings of this study.

**Initial observation and alignment structure**

In most PLR-containing reads (Supplementary Table 4), basecalls are aligned exclusively to the forward template sequence. Only a subset of reads exhibited the expected template-peptide-threading configuration (Supplementary Table 10). Initial inspection suggested that peptide-pore interactions reduced basecalling accuracy in the downstream threading region. However, further analysis revealed that in most reads the downstream segment corresponded to the reverse complement of the template rather than the threading sequence.

*De novo* multiple sequence alignment using MAFFT[1] revealed a bipartite structure consisting of a forward-aligned template segment followed by its reverse complement. To validate this observation, we performed reference-based alignment using minimap2 against a synthetic reference consisting of the template concatenated to its reverse complement (Supplementary Table 9). In these alignments, reads typically map from positions ~1-60 of the forward template and subsequently transition to the RC strand near the same position and extending toward position 1 (approximately positions 80-120 in the concatenated reference). RC segments exhibited near-complete sequence identity and consistently high basecalling quality ($Q > 20$). Transition points occurred within a narrow window of ±5 bases across independent runs (Supplementary Fig. 22).

**Artefacts in reads aligning exclusively to threading DNA**

A similar sequencing artefact was observed for reads that align exclusively to the threading DNA (Supplementary Figure 16). These reads originate from sequencing runs containing a mixture of template, threading, and peptide constructs, but represent a subset of reads in which only the threading sequence is aligned. We interpret these reads as originating from molecules consisting of threading DNA without an attached peptide.

In these cases the forward alignment spanned position 1 to 20, followed by a reverse-complement alignment from ~30 back to position 1. Unlike PLR-containing reads, these events showed no current drop between segments. Signal traces were continuous and of high quality. This indicates that the phenomenon is independent of the peptide insertion.

**Reproducibility and quantitative distribution**

Across sequencing runs, we quantified the fraction of reads aligning exclusively to the forward template, the expected template-threading configuration, and the template-RC-template configuration (Supplementary Table 10). Comparable proportions were observed across experiments.

**Raw signal characteristics**
Inspection of raw ionic current traces using the input POD5 files and the Bulk fast5 files revealed no signal interruption or voltage reversal at the forward-to-RC transition. The current trace remained continuous and motor stepping appeared uninterrupted, with basecalling quality consistently above Q20. These observations indicate that the RC segment arises during a single continuous translocation event rather than from pore re-engagement or sequencing restart.

**Experimental controls**
Several control experiments were performed to investigate potential causes of the RC configuration.

*Library preparation controls*
To assess whether individual library preparation steps contributed to the RC read formation, we systematically modified the protocol. Omitting the nick repair step during library preparation did not reduce the frequency of RC reads, indicating that nick repair is unlikely to underlie this configuration. In a separate control experiment, the adapter ligation step was omitted entirely. Under these conditions, no signals aligning to the control DNA or template sequence were detected, confirming that sequencing events depend on successful adapter ligation and demonstrating that ligation proceeds efficiently for the single-stranded constructs used in this study.

*Construct design controls*
Early experiments used constructs containing a double-stranded template region generated by annealing a complementary strand prior to adapter ligation. Reverse-complement chimeras were observed in both double-stranded and single-stranded constructs at comparable frequencies, indicating that the phenomenon is independent of template strand configuration during library preparation. However, single-stranded constructs yielded substantially higher numbers of detectable PLRs and were therefore used in subsequent experiments (Supplementary Table 12).

*Peptide-free control experiments*
Sequencing of template DNA without peptide or threading DNA produced similar forward-RC chimeras. These reads showed continuous signal without the low-amplitude PLR segment, demonstrating that the phenomenon does not depend on peptide linkage (Supplementary Fig. 19).

*Computational validation*
The RC configuration was confirmed using multiple basecalling models and versions (SUP v5.2.0, v5.0.0; Dorado v1.0.0 and v0.9.0) as well as independent alignment approaches (minimap2 and MAFFT), supporting that it is not a computational artefact.

**Proposed mechanism**
The observed RC configuration is best explained by strand extension initiated through internal priming during library preparation, a mechanism supported by personal communication with R&D scientists at New England Biolab and application scientist at Oxford Nanopore Technologies, who reported that strand-extension polymerase activity is present within the end-repair enzyme mixture. End-repair reactions are known to enable extension from

transiently formed 3′ self-complementary structures, such as hairpins [2,3]. In single-stranded DNA, intramolecular base pairing can generate short hairpin structures that provide a priming substrate for polymerase-mediated extension, resulting in the synthesis of a covalently linked reverse-complement segment. This mechanism is consistent with established snap-back and fold-back DNA synthesis phenomena, as well as known artefact formation during enzymatic library preparation workflows [4,5].

This interpretation is supported by several observations. The RC segments exhibit high basecalling accuracy, transition points are highly reproducible, and the phenomenon occurs independently of peptide presence. In addition, secondary structure prediction using UNAFold identified stable hairpin structures near template position ~60 and threading position ~20, coinciding with the observed forward-to-RC transition sites (Supplementary Fig. 20–23). Together, these findings are consistent with templated strand extension initiated at preferred secondary structure sites.

**Alternative explanations for RC chimeras**

Alternative explanations are less consistent with the observed data. Voltage reversal or re-reading events would be expected to introduce signal discontinuities or motor dissociation, neither of which are observed. Purely structural fold-back translocation without strand synthesis is insufficient to explain the high sequence accuracy of the RC segment. Computational artefacts are likewise unlikely, as the phenomenon is consistently observed across multiple basecalling models and alignment strategies.

**Impact on peptide-linker regions and classification**

The presence of RC segments does not affect peptide signal analysis. The PLR consistently precedes the forward-to-RC transition and exhibits highly reproducible signal characteristics independent of alignment configuration. To this end, PLRs were extracted from reads aligning either to the canonical template-threading configuration or the template-RC-template configuration. Signal similarity quantified using dynamic time warping and feature extraction showed comparable within-class distance distributions across alignment categories (Supplementary Fig. 17), indicating that peptide-associated signal signatures are preserved.

To further assess robustness, the classifier was trained and validated exclusively on PLRs from template-RC-template reads and subsequently evaluated on reads aligning to the canonical template-threading configuration. Despite the fewer number of template-threading reads, relative classification performance across peptides remained comparable (Supplementary Fig. 18).

**Conclusion**

Chimeric forward-reverse-complement template reads occur reproducibly in a consistent fraction of sequencing events across independent runs. These reads arise during continuous translocation, are independent of peptide presence and construct configuration, and persist across library preparation variations and computational pipelines. Importantly, this phenomenon does not interfere with peptide detection or classification, and all central conclusions of this study remain unaffected.

**Supplementary Note 02 | Signal refinement of nanopore sequencing signals**

Basecaller-emitted move-table (mv / move tags) information provides a quick mapping from called bases to positions in the raw current trace. However, the initial move table represents only a coarse mapping. To obtain reliable, per-base signal assignments that are comparable across reads we apply a post-basecalling signal-refinement step (re-squiggling) using remora utilities (v0.6.0). This step improves segmentation of the continuous current trace into DNA base-associated segments and performs per-read scaling and shift corrections, thereby removing baseline and scaling differences between runs and reads.

The presence of peptides introduces signal regions whose current characteristics differ from canonical DNA k-mer expectations. The PLR can generate current levels outside the range typically modeled by DNA-only pore models, influencing signal-to-reference alignment and scaling behavior during refinement. To ensure stable refinement and normalization across reads, we adjusted several refinement parameters:

- **do_fix_gauge = True**: constrains the refined signal distribution to a mean near 0 and a robustly estimated standard deviation near 1 using median- and MAD-based normalization, improving consistency across reads.
- **do_rough_rescale = False**: disables approximate scaling updates derived from the move table to avoid propagating local scaling biases from non-canonical signal regions into neighboring DNA regions
- **scale_iters = 0**: performs a single precise signal-to-sequence refinement pass without iterative re-scaling, improving stability in regions with non-canonical current behavior.

**Supplementary Note 03 | PLR segmentation**

PLRs were identified in individual nanopore reads using a signal-based detection algorithm designed to detect characteristic current drops associated with peptide translocation while remaining robust to signal noise, baseline drift, and minor alignment inaccuracies. The algorithm combines alignment-derived priors with signal-level heuristics to identify contiguous regions exhibiting peptide-like current signatures.

1. **Signal extraction and mapping**
   Raw current signals were obtained from POD5 reads via remora's io.Read.from_pod5_and_alignment, guided by the corresponding SAM alignment. Signal-to-sequence mapping was refined using Remora's signal mapping refinement procedure to improve correspondence between nucleotide positions and signal indices. The refined mapping provides a query_to_signal index that enables conversion between sequence positions and raw signal coordinates.
2. **Alignment-based search window**
   To localize the DNA-peptide transition, an anchor reference position corresponding to the end of the template DNA was defined. If this position was not aligned in a read, the nearest preceding aligned reference coordinate was used.
   The anchor was converted to signal coordinates using the refined sequence-to-signal mapping. A symmetric search window centered on this position was then defined, spanning ±70 bases in sequence space. This window was converted to signal coordinates using the median stride of the sequence-to-signal mapping to account for variability in signal sampling density.
3. **DNA baseline estimation**
   Within the search window, the DNA baseline current was estimated from the upstream portion of the signal corresponding to DNA. The baseline was defined as the median current of the left fraction of the window (25% of the window length, up to a maximum of 1000 samples), providing a robust estimate of the DNA current level for each read.
4. **Detection of candidate regions**

Candidate peptide regions were detected as sustained decreases in current relative to the DNA baseline. For each signal sample, the current drop relative to the baseline was computed. To capture drops occurring at different temporal scales, this signal was smoothed using uniform filters with window sizes of 15 and 35 samples. The maximum of the two smoothed signals was used as a combined drop feature. Samples exceeding a drop threshold were marked as candidate regions. The threshold corresponded to 45% of the minimum pA-drop criterion (with a minimum absolute threshold of 5 pA).

5. **Candidate scoring and selection**
   For each candidate region, the region length and signal drop relative to the DNA baseline were computed. The signal drop was defined as the difference between the DNA baseline and the 25th percentile current within the region. In addition, the deviation from the expected move-level current predicted by the Remora signal model was calculated.

   Candidates were ranked using a combined score defined as the sum of the signal drop and the move-level deviation. Regions were required to satisfy the following criteria:

   - minimum length ≥ 60 signal samples
   - pA drop ≥ 75 pA
   - move-level deviation ≥ 15 pA
   - distance to the DNA-aligned region ≤ 500 signal samples

   The highest-scoring candidate satisfying these criteria was selected as the PLR for the read.

6. **Boundary refinement**
   PLR boundaries were adjusted to the nearest indices in the sequence-to-signal mapping to maintain alignment consistency. Reads were excluded if the detected region occurred within the final 10% of the signal trace or if the surrounding signal did not resemble typical DNA current levels (e.g., excessive low-current or non-finite values outside the PLR).

**Supplementary Note 04 | Feature extraction overview**

To comprehensively characterize nanopore peptide signal traces, we first constructed a feature library capturing complementary aspects of signal amplitude, distributional shape, and temporal dynamics of the nanopore current signal. For each read, an initial set of 152 time-series features was computed.

The full feature set comprised several categories:

- Descriptive statistics: central tendency, dispersion, and robust quantiles (e.g., mean, median, IQR, Q10–Q90, trimean, midhinge).
- Distributional shape: skewness, kurtosis, MAD-based measures, and coefficient of variation.
- Temporal dynamics: first- and second-order derivative statistics, including slope and curvature descriptors.
- Local structure: rolling-window slope statistics (window size: 100 samples).
- Extrema-based features: peak/trough counts, prominence, and spacing statistics.

- Global signal properties: energy, RMS, AUC, and detrended variance.
- Transient drop features: magnitude, duration, and position of dominant downward transitions.
- Catch22 features: 22 canonical time-series descriptors capturing nonlinear dynamics [6]

To reduce redundancy and improve generalization, feature selection was performed exclusively on the training set using a balanced subset of 10,000 reads per class. The procedure included variance filtering, removal of highly correlated features ($|r| > 0.95$), and ranking based on a combination of mutual information and gradient-boosted tree importance.

Model performance was evaluated using progressively smaller subsets of ranked features. A final set of 50 features was selected at the point of performance saturation and used for all downstream classification analyses (Supplementary Table 12). This reduction yielded a compact representation while preserving predictive performance and reducing redundancy.

### Supplementary Tables

**Supplementary Table 1 | Sequences of DNA oligonucleotides and the Peptide-Oligonucleotide Construct.**

| Name | Sequence [5’ -> 3’] |
|---|---|
| Template DNA | tta ctg aag tct cac gtg cct ggt ata tta gcg tcc act ctc act atc gga ttc tac atc ggt cgt agc c-Maleimid |
| Threading DNA | DBCO-PEG-PEG-PEG-PEG-agc tga tcg atc gta gct agc tag cat gct caa aaa aaa aaa aaa aaa aaa aaa aaa aaa |
| Peptide-Oligonucleotide Construct | tta ctg aag tct cac gtg cct ggt ata tta gcg tcc act ctc act atc gga ttc tac atc ggt cgt agc c -3`-Maleimid-[PEPTIDE]-PEG-PEG-PEG-PEG-Triazid-DBCO-PEG-PEG-PEG-PEG-5`-agc tga tcg atc gta gct agc tag cat gct caa aaa aaa aaa aaa aaa aaa aaa aaa aaa |

**Supplementary Table 2 | Sequences and purification characteristics of the analyzed peptides.**
Peptide sequences are presented from the N-terminus to the C-terminus, corresponding to the direction of translocation through the nanopore.

| Name | Sequence [N -> C] | Modification [N-term] | Modification [C-term] | Purity | Vendor |
|---|---|---|---|---|---|
| ßCAT | YLDSGIHSGAC | {N3-PEG4} | - | >90% | JPT Peptide Technologies GmbH |
| ßCAT-D | YLDSDIHSGAC | {N3-PEG4} | - | ≥90% | GenScript Biotech |
| ßCAT-W | YLDSWIHSGAC | {N3-PEG4} | - | ≥90% | GenScript Biotech |
| ßCAT-GG | YLDSGGHSGAC | {N3-PEG4} | - | ≥90% | GenScript Biotech |
| ßCAT-L | YLDSLIHSGAC | {N3-PEG4} | - | ≥90% | GenScript Biotech |
| ßCAT-WW | YLDSWWHSGAC | {N3-PEG4} | - | ≥90% | GenScript Biotech |
| ßCAT-WWW | YLDSWWWHSGAC | {N3-PEG4} | - | ≥90% | GenScript Biotech |
| ßCAR3 | IMDRTPEKLC | {N3-PEG4} | - | >90% | JPT Peptide |

| | | | | | |
|---|---|---|---|---|---|
| | | | | | Technologies GmbH |
| ßCAT-30 | YLDSGIHSGACKTGKHGEGCEAVKLQRDLC | {N3-PEG4} | - | ≥90% | GenScript Biotech |
| ßCAT-35 | YLDSGIHSGACKTGKHGEGCEAVKLQRDLGCDLQC | {N3-PEG4} | - | ≥90% | GenScript Biotech |
| CHG-negG | YEYEYEGEYEYEC | {N3-PEG4} | - | crude | GenScript Biotech |
| CHG-posD | RKHGRKWHDKRKC | {N3-PEG4} | - | ≥90% | GenScript Biotech |
| CHG-posL | RKHGRKWHLKRKC | {N3-PEG4} | - | ≥90% | GenScript Biotech |
| SZ-small | GSGAGSSGGSIGGRC | {N3-PEG4} | - | ≥90% | GenScript Biotech |
| SZ-large | GFLFPEHTYFFRC | {N3-PEG4} | - | ≥90% | GenScript Biotech |
| βCAT-Hphil | YLDSGIHSGAKDKKC | {N3-PEG4} | - | ≥90% | GenScript Biotech |
| βCAT-Hmod | YLDSGIHSGALKGQC | {N3-PEG4} | - | ≥90% | GenScript Biotech |
| βCAT-Hphob | YLDSGIHSGAKKKAC | {N3-PEG4} | - | ≥90% | GenScript Biotech |
| βCAT-ins1 | VVVVGYLDSGIHSGAC | {N3-PEG4} | - | crude | GenScript Biotech |
| βCAT-ins2 | YLDSGIHSGAGVVVVC | {N3-PEG4} | - | crude | GenScript Biotech |

**Supplementary Table 3 | Physicochemical properties and characteristics of sequenced peptides.**

Peptide sequences are listed from the N-terminus to the C-terminus, corresponding to the direction of translocation through the nanopore. Net charges at pH 7.0 were calculated using a Henderson-Hasselbalch model with Bjellqvist $pK_a$ values as implemented in Biopython (v1.86), with the N-terminal charge contribution omitted to account for N-terminal azide modification. Molecular weights (average mass) were adjusted accordingly for the N-terminal modification. Hydrophobicity values (GRAVY index; Kyte-Doolittle scale) were calculated using Biopython ProtParam.

| Name | Sequence (N->C) | Length (aa) | Net charge (pH 7.0; N-term blocked) | Mass (Da, average) | GRAVY index (Kyte-Doolittle) |
|---|---|---|---|---|---|
| ßCAT | YLDSGIHSGAC | 11 | -1.92 | 1163.22 | 0.200 |
| ßCAT-D | YLDSDIHSGAC | 11 | -2.92 | 1221.26 | -0.082 |
| ßCAT-W | YLDSWIHSGAC | 11 | -1.92 | 1292.38 | 0.155 |
| ßCAT-GG | YLDSGGHSGAC | 11 | -1.92 | 1107.11 | -0.245 |
| ßCAT-L | YLDSLIHSGAC | 11 | -1.92 | 1219.33 | 0.582 |
| ßCAT-WW | YLDSWWHSGAC | 11 | -1.92 | 1365.43 | -0.336 |
| ßCAT-WWW | YLDSWWWHSGAC | 12 | -1.92 | 1551.64 | -0.383 |
| BCAR3 | IMDRTPEKLC | 10 | -1.01 | 1246.46 | -0.500 |
| ßCAT-30 | YLDSGIHSGACKTGKHGEGCEAVKLQRDLC | 30 | -0.85 | 3217.57 | -0.483 |
| ßCAT-35 | YLDSGIHSGACKTGKHGEGCEAVKLQRDLGCDLQC | 35 | -1.86 | 3734.14 | -0.446 |
| CHG-negG | YEYEYEGEYEYEC | 13 | -7.00 | 1809.77 | -1.954 |
| CHG-posD | RKHGRKWHDKRKC | 13 | +5.16 | 1776.04 | -2.908 |
| CHG-posL | RKHGRKWHLKRKC | 13 | +6.16 | 1774.11 | -2.346 |
| SZ-small | GSGAGSSGGSIGGRC | 15 | -0.01 | 1250.26 | -0.113 |
| SZ-large | GFLFPEHTYFFRC | 13 | -0.92 | 1704.91 | 0.177 |

| | | | | | |
|---|---|---|---|---|---|
| βCAT-Hphil | YLDSGIHSGAKDKKC | 15 | +0.08 | 1662.82 | -0.867 |
| βCAT-Hmod | YLDSGIHSGALKGQC | 15 | -0.92 | 1589.73 | -0.120 |
| βCAT-Hphob | YLDSGIHSGAKKKAC | 15 | +1.07 | 1618.82 | -0.513 |
| βCAT-ins1 | VVVVGYLDSGIHSGAC | 16 | -1.92 | 1616.8 | 1.163 |
| βCAT-ins2 | YLDSGIHSGAGVVVVC | 16 | -1.92 | 1616.8 | 1.162 |

**Supplementary Table 4 | Summary statistics for MinION nanopore sequencing runs.** For each sequencing run, the table reports the run number and identifier, the target peptide name, the number of basecalled reads, the number of reads aligned to the reference construct, and the number of extracted PLRs together with the percentage relative to the number of aligned reads. The final column reports the downstream usage of PLRs from each sequencing runs: Runs labeled *Train-Valid* were included in model training and validation; Runs labeled *Test* were held out entirely and used exclusively as an independent test set; Runs marked with *"-"* were excluded from all analyses due to insufficient yield or quality. Runs labeled as "DNA control" correspond to experiments in which only template DNA was included in the sequencing mixture, without threading strands or DNA-peptide conjugates.

| Run # | Run ID | Peptide name | Basecalled reads (n) | Aligned reads to template (n) | PLRs (n, % of aligned) | Data split |
|---|---|---|---|---|---|---|
| 1 | FAV99375_a2f3daba_f5d5051e | DNA control (only template) | 756,418 | 45,096 | 69 (0.15%) | - |
| 2 | FAV99258_6691a984_21de4d21 | ßCAT | 157,948 | 6,094 | 16 (0.26%) | Train-Valid |
| 3 | FAV99364_e5b921e6_20d8415a | BCAR3 | 429,790 | 64,596 | 13,296 (20.58%) | Train-Valid |
| 4 | FBA38751_61ba46ff_a079865e | ßCAT | 463,413 | 21,413 | 12,099 (56.50%) | Test |
| 5 | FBA38661_d8d1d6fb_cdc15b3a | ßCAT | 148,906 | 32,016 | 16,804 (52.49%) | Train-Valid |

| | | | | | | |
|---|---|---|---|---|---|---|
| 6 | FBA29925_da2ca5cc_9c225447 | ßCAT | 164,408 | 40,575 | 24,990 (61.59%) | Train-Valid |
| 7 | FBA36690_11283c27_1d68b023 | ßCAT-30 | 1,327,915 | 14,162 | 1,031 (7.28%) | Train-Valid |
| 8 | FBA36633_3e637743_0c4ddd7f | ßCAT-35 | 399,595 | 26,838 | 586 (2.18%) | Train-Valid |
| 9 | FBA36696_12b23436_aa892ef3 | ßCAT-D | 460,056 | 46,735 | 24,298 (51.99%) | Test |
| 10 | FBA38740_4bdb99f1_8ee27ae2 | ßCAT-W | 724,935 | 19,717 | 13,561 (68.78%) | Train-Valid |
| 11 | FBA30048_b5ceaa16_db65ef03 | ßCAT-L | 1,355,013 | 17,723 | 12,235 (69.03%) | Train-Valid |
| 12 | FBA36641_f625ddb5_1f8e472d | SZ-small | 1,438,486 | 92,340 | 14,282 (15.47%) | Train-Valid |
| 13 | FBA38709_9268d485_1cbb4986 | SZ-large | 1,009,026 | 1,450 | 577 (39.79%) | Test |
| 14 | FBA28482_67b2332e_6be9a788 | ßCAT-Hmod | 1,250,192 | 18,822 | 4,943 (26.26%) | Train-Valid |
| 15 | FBA30116_e02496d5_ad097796 | ßCAT-Hphob | 1,793,882 | 23,538 | 5,187 (22.04%) | Train-Valid |
| 16 | FBA27675_a78a15c7_5bf1930c | ßCAT-Hphil | 1,260,917 | 154,981 | 39,964 (25.79%) | Train-Valid |
| 17 | FBA27715_f3322461_d486bb6d | CHG-negG | 512,150 | 5,213 | 2,472 (47.42%) | Train-Valid |
| 18 | FBA29916_cac7b305_26891754 | CHG-posD | 528,805 | 3,448 | 41 (1.19%) | - |
| 19 | FBA27966_5469d454_4d69982d | CHG-posL | 827,114 | 4,446 | 63 (1.41%) | - |

| | | | | | | |
|---|---|---|---|---|---|---|
| 20 | FBA28478_6423cf66_bb59bd64 | ßCAT-ins-1 | 1,476,056 | 4,960 | 1,031 (20.79%) | Train-Valid |
| 21 | FBA27518_fd8611d5_29122d7e | ßCAT-ins-2 | 565,648 | 3,668 | 2,052 (55.94%) | Train-Valid |
| 22 | FBA27515_98e7cb06_6887d93b | ßCAT-GG | 897,994 | 120,419 | 84,482 (70.16%) | Train-Valid |
| 23 | FBA27474_e80cd68c_59548e22 | ßCAT-WW | 806,300 | 145,534 | 108,556 (74.59%) | Train-Valid |
| 24 | FBA29994_e0c5fddc_e4e5c004 | ßCAT-WWW | 809,665 | 74,858 | 55,722 (74.44%) | Train-Valid |
| 25 | FBA29985_b5dcf3a7_8385330d | SZ-large | 299,414 | 3,610 | 1,918 (53.13%) | Train-Valid |
| 26 | FBA38608_530747f8_e1cc8434 | ßCATWW | 917,902 | 60,077 | 43,434 (72.30%) | Test |
| 28 | FBA32908_2198cdfd_925f5b24 | Mixture 1 | 29,480 | 3,623 | 2,296 (63.37%) | Test |
| 29 | FBA27747_42ec133f_e0f5aa10 | Mixture 2 | 1,194,106 | 32,712 | 8,683 (26.54%) | Test |
| 30 | FBC91212_096c66d1_96a02039 | ßCAT-D | 912,839 | 181,132 | 127,631 (70.46%) | Train-Valid |
| 31 | FBA27571_cdeed201_769ebcc8 | Mixture 1 | 848,345 | 151,644 | 99,658 (65.72%) | Test |

**Supplementary Table 5 | Performance metrics across the full dataset for different models.**
Accuracy, weighted F1, and macro-averaged F1 scores are reported as percentages (0-100) for both the validation and test sets. InceptionTime and MiniRocket were implemented using the aeon library. For MiniRocket, default parameters were used, with the number of kernels set to 1,000.

| Model | Validation | | | Test | | |
|---|---|---|---|---|---|---|
| | Accuracy | Weighted F1-score | Macro F1-score | Accuracy | Weighted F1-score | Macro F1-score |
| InceptionTime | 83.07 | 84.41 | 63.74 | 78.16 | 85.84 | 74.00 |
| MiniRocket | 71.12 | 66.42 | 33.74 | 77.14 | 79.14 | 50.83 |
| features -LGBM | 66.19 | 67.90 | 49.07 | 58.99 | 70.92 | 59.13 |

**Supplementary Table 6 | Per-class validation performance metrics for the LightGBM and InceptionTime model.**
F1 scores are reported as percentages (0-100).

| Peptide | F1 LightGBM | F1 InceptionTime |
|---|---|---|
| BCAR3 | 49.21 | 71.74 |
| CHGnegG | 32.99 | 54.90 |
| SZlarge | 30.59 | 34.39 |
| SZsmall | 71.91 | 87.93 |
| ßCAT | 44.18 | 72.47 |
| ßCAT30 | 22.22 | 23.65 |
| ßCAT35 | 17.24 | 16.80 |
| ßCATD | 75.72 | 90.00 |
| ßCATGG | 73.86 | 90.05 |
| ßCATL | 35.15 | 73.04 |
| ßCATW | 34.71 | 59.58 |
| ßCATWW | 68.22 | 84.66 |
| ßCATWWW | 69.96 | 85.71 |
| βCATHmod | 51.22 | 68.44 |
| βCATHphil | 87.69 | 93.27 |
| βCATHphob | 77.61 | 73.31 |
| βCATins1 | 25.57 | 49.58 |

| | | |
|---|---|---|
| βCATins2 | 15.24 | 17.88 |

**Supplementary Table 7 | Per-class test performance metrics for the LightGBM and InceptionTime model.**
F1 scores are reported as percentages (0-100).

| Peptide | F1 LightGBM | F1 InceptionTime |
|---|---|---|
| SZlarge | 32.80 | 42.16 |
| ßCAT | 53.00 | 77.15 |
| ßCATD | 78.58 | 90.40 |
| ßCATWW | 72.14 | 86.28 |

**Supplementary Table 8 | Validation performance of the InceptionTime model with varying numbers of included peptides.**
Accuracy, weighted F1, and macro-averaged F1 scores are reported as percentages (0-100) for peptide sets ranging from 2 to 18.

| Set size | Accuracy | Weighted F1 | Macro F1 |
|---|---|---|---|
| 2 (ßCAT, ßCAT-W) | 98.57 | 98.58 | 98.09 |
| 4 (ßCAT, ßCAT-W, ßCAT-L, ßCAT-D) | 92.28 | 92.39 | 88.93 |
| 7 (ßCAT, ßCAT-W, ßCAT-L, ßCAT-D, ßCAT-GG, ßCAT-WW, ßCAT-WWW) | 89.96 | 90.19 | 84.89 |
| 18 | 83.06 | 84.41 | 63.74 |

**Supplementary Table 9 | Sequences of DNA oligonucleotides used for minimap2 alignment**

| Name | Sequence |
|---|---|
| Template | TTACTGAAGTCTCACGTGCCTGGTATATTAGCGTCCACTCTCACTATCGGATTCTACATCGGTCGTAGCC |
| Threading | AGCTGATCGATCGTAGCTAGCTAGCATGCTCAAAAAAAAAAAAAAAAAAAAAAAAAAAA |
| Template-Threading | TTACTGAAGTCTCACGTGCCTGGTATATTAGCGTCCACTCTCACTATCGGATTCTACATCGGTCGTAGCCAGCCAGCTGATCGATCGTAGCTAGCTAGCATGCTCAAAAAAAAAAAAAAAAAAAAAAAAAAAAA |
| Template-RC-Template | TTACTGAAGTCTCACGTGCCTGGTATATTAGCGTC |

| | |
|---|---|
| | CACTCTCACTATCGGATTCTACATCGGTCGTAGC<br>CGGCTACGACCGATGTAGAATCCGATAGTGAGAG<br>TGGACGCTAATATACCAGGCACGTGAGACTTCAG<br>TAA |
| Threading-RC-Threading | AGCTGATCGATCGTAGCTAGCTAGCATGCTCAAA<br>AAAAAAAAAAAAAAAAAAAAAAAAAATTTTTTTTTT<br>TTTTTTTTTTTTTTTTTTTGAGCATGCTAGCTAGCT<br>ACGATCGATCAGCT |

**Supplementary Table 10 | Alignment statistics using different reference sequences for nanopore sequencing runs.**
For each run, the table reports the run number, target peptide, and the number of reads aligned to three reference sequences: template, template-threading, and template-RC-template. Only reads from the positive strand were considered, to avoid reverse-complement alignments of the template to the second part of the reference. Template alignments were counted only if they covered at least position 59. For template-threading and template-RC-template, only full-length alignments were included, which were defined as reads starting before position 30 and extending to at least position 110. This criterion ensures that partial alignments overlapping the standard template region do not artificially inflate read counts.

| Run Number | Peptide name | # of aligned to template | # of aligned to template-threading | # of aligned to template-RC-template |
|---|---|---|---|---|
| 1 | - (only DNA) | 27,153 | 27 | 12,771 |
| 2 | ßCAT | 3,030 | 7 | 1,975 |
| 3 | BCAR3 | 38,587 | 796 | 20,557 |
| 4 | ßCAT | 13,225 | 1,390 | 6,623 |
| 5 | ßCAT | 18,006 | 459 | 13,229 |
| 6 | ßCAT | 26,023 | 930 | 16,321 |
| 8 | ßCAT-30 | 9,716 | 221 | 3,211 |
| 9 | ßCAT-35 | 19,210 | 195 | 6,271 |
| 10 | ßCAT-D | 31,276 | 1,328 | 16,222 |
| 11 | ßCAT-W | 11,479 | 183 | 7,647 |
| 12 | ßCAT-L | 10,469 | 343 | 6,759 |
| 13 | SZ-small | 58,658 | 648 | 37,905 |
| 14 | SZ-large | 756 | 534 | 378 |
| 15 | ßCAT-Hmod | 11,775 | 983 | 6,613 |
| 16 | ßCAT-Hphob | 14,524 | 1,133 | 7,343 |

| | | | | |
|---|---|---|---|---|
| 17 | ßCAT-Hphil | 89,675 | 2,190 | 64,712 |
| 18 | CHG-negG | 2,892 | 668 | 2,037 |
| 19 | CHG-posD | 2,015 | 698 | 188 |
| 20 | CHG-posL | 2,430 | 1,400 | 274 |
| 21 | ßCAT-ins-1 | 3,022 | 1,572 | 1,768 |
| 22 | ßCAT-ins-2 | 2,135 | 1,249 | 1,170 |
| 23 | ßCAT-GG | 73,272 | 2,946 | 50,334 |
| 24 | ßCAT-WW | 77,203 | 3,471 | 60,262 |
| 25 | ßCAT-WWW | 34,050 | 1,884 | 29,299 |
| 26 | SZ-large | 1,781 | 2,376 | 1,217 |
| 27 | ßCATWW | 30,992 | 1,212 | 23,201 |
| 29 | Mixture 1 | 2,132 | 43 | 1,513 |
| 30 | Mixture 2 | 20,330 | 1,409 | 12,654 |
| 31 | ßCAT-D | 121,116 | 2,012 | 73,661 |
| 32 | Mixture 1 | 96,949 | 3,176 | 59,477 |

**Supplementary Table 11 | Comparison of βCAT single-strand and double-strand variants.**
For each sequencing run, the table reports the run number and identifier, the target peptide, the number of basecalled reads, the number of reads aligned to the reference construct, and the number of extracted PLRs. The final column shows the number of PLRs along with the percentage relative to aligned reads.

| Name | # of basecalls | # of alignments to template | # of detected PLRs |
|---|---|---|---|
| ßCAT-ss-01 | 463,413 | 21,413 | 12,099 (56.50%) |
| ßCAT-ss-02 | 148,906 | 32,016 | 16,804 (52.49%) |
| ßCAT-ss-03 | 164,408 | 40,575 | 24,990 (61.59%) |
| ßCAT-ds-01 | 380,851 | 227,657 | 553 (0.24%) |
| ßCAT-ds-02 | 253,795 | 130,148 | 493 (0.38%) |
| ßCAT-ds-03 | 437,556 | 214,087 | 643 (0.30%) |

**Supplementary Table 12 | Top 50 selected features used for classification.**

| Feature | Description |
|---|---|
| min_step_height | Minimum detected step amplitude |
| rolling_slope_q25 | 25th percentile of rolling slopes |
| min_slope | Minimum first-derivative value |
| skew | Signal distribution skewness |
| c22_DN_HistogramMode_10 | Mode of histogram (10 bins) |
| c22_DN_OutlierInclude_p_001_mdrmd | Outlier structure statistic |
| cross_rate_q25 | 25th percentile zero-crossing rate |
| c22_DN_HistogramMode_5 | Mode of histogram (5 bins) |
| drop_depth | Magnitude of largest signal drop |
| drop_auc_norm | Normalized area of drop region |
| slope_q60 | 60th percentile of signal slope |
| drop_auc | Area under drop segment |
| max | Maximum signal amplitude |
| slope_q75 | 75th percentile of slope |
| mean_peak_prom | Mean prominence of peaks |
| drop_idx | Position of largest drop |
| c22_CO_trev_1_num | Time-reversibility statistic |
| drop_post_min | Minimum value after drop |
| drop_duration | Duration of drop event |
| median_rolling_slope | Median rolling-window slope |
| kurtosis | Signal distribution kurtosis |
| c22_SC_FluctAnal_2_dfa_50_1_2_logi_prop_r1 | Detrended fluctuation scaling |
| total_variation | Sum of absolute slope changes |
| drop_pre_mean | Mean signal before drop |
| c22_DN_OutlierInclude_n_001_mdrmd | Negative outlier statistic |
| slope_q90 | 90th percentile of slope |
| mean_step_height | Mean detected step amplitude |

| | |
|---|---|
| MAD_over_IQR | Robust variability ratio |
| idl_60_40 | Inter-decile level difference |
| c22_CO_HistogramAMI_even_2_5 | Histogram-based auto mutual information |
| c22_SC_FluctAnal_2_rsrangefit_50_1_logi_prop_r1 | Rescaled range fluctuation statistic |
| max_step_height | Maximum detected step amplitude |
| c22_MD_hrv_classic_pnn40 | Proportion of large differences |
| c22_SB_MotifThree_quantile_hh | Motif distribution statistic |
| min_rolling_slope | Minimum rolling slope |
| q60 | 60th percentile signal value |
| trimmed_MAD | Trimmed median absolute deviation |
| median | Median signal value |
| cross_rate_q10 | 10th percentile crossing rate |
| cross_rate_q50 | Median crossing rate |
| median_slope | Median signal slope |
| max_rolling_slope | Maximum rolling slope |
| rolling_slope_q60 | 60th percentile rolling slope |
| slope_q10 | 10th percentile slope |
| range | Signal value range |
| mean_slope | Mean signal slope |
| q90 | 90th percentile signal value |
| q75 | 75th percentile signal value |
| c22_FC_LocalSimple_mean1_tauresrat | Autocorrelation decay ratio |
| mean_peak_dist | Mean distance between peaks |

### Supplementary Figures

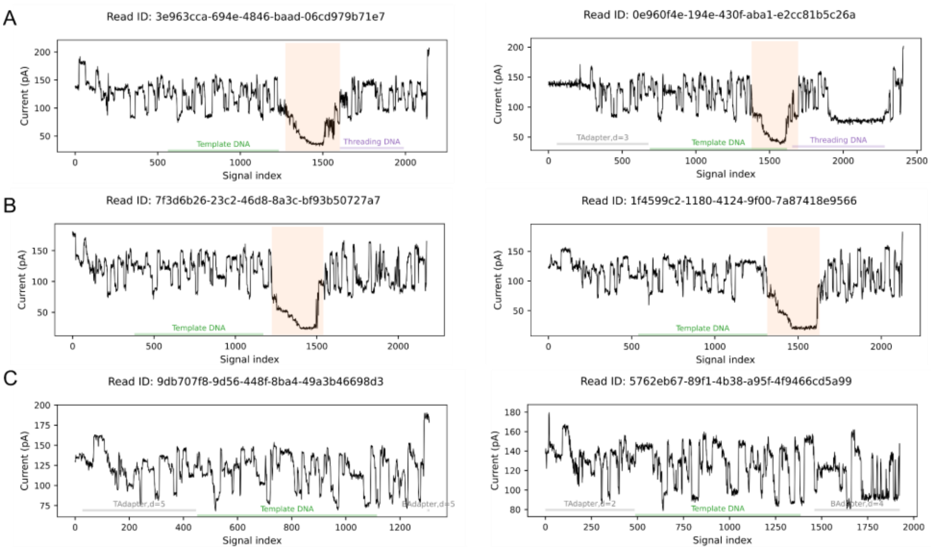


**Supplementary Figure 1 | Segmentation of βCAT signals.**
**(A)** Representative signals with a detected peptide-linked region (PLR), located between aligned template and threading DNA segments. **(B)** Representative signals with a detected PLR, but with alignment only to the template DNA. **(C)** Representative signals with template DNA alignment but no detectable PLR.

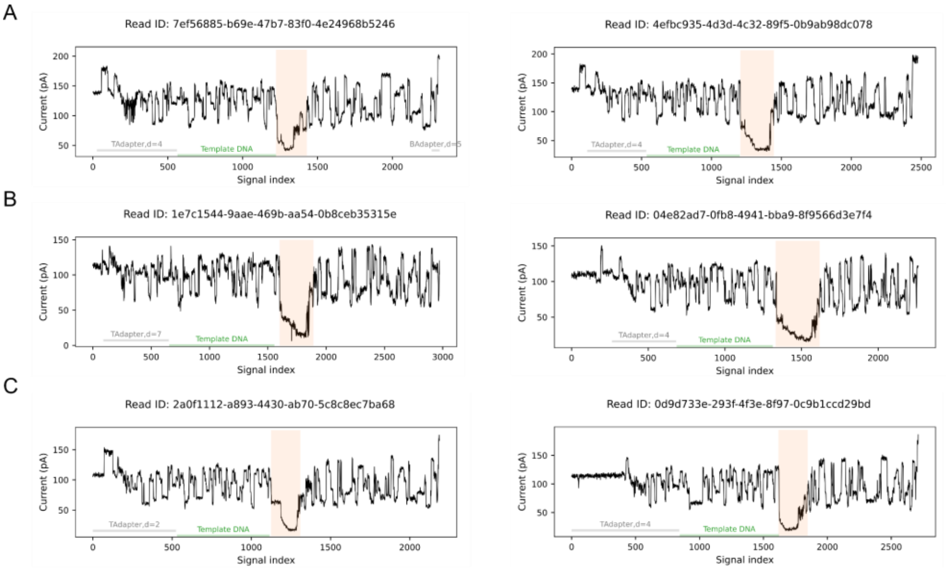


**Supplementary Figure 2 | Reproducibility of PLR signals.**
Representative βCAT signals from replicate runs: rep1 **(A)**, rep2 **(B)**, and rep3 **(C)**. Measurements across independent flow cells show consistent PLR signal characteristics, including similar amplitude profiles.

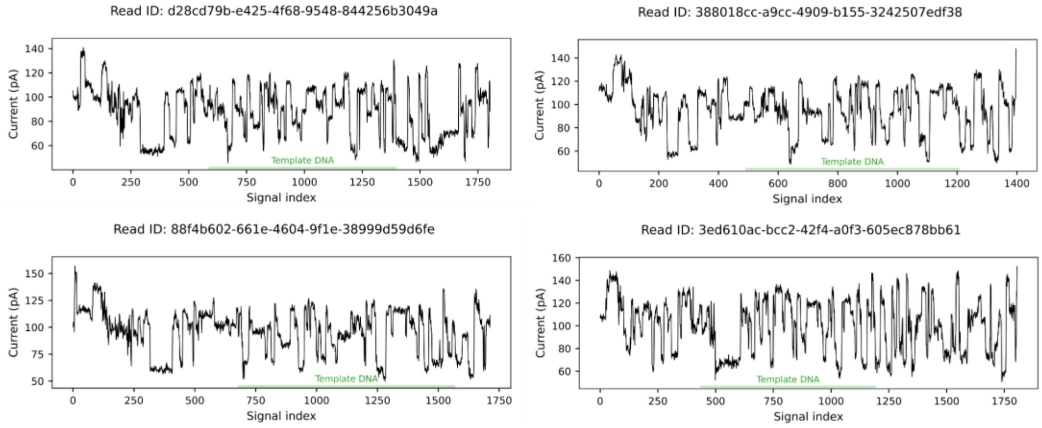


**Supplementary Figure 3 | Control experiments with template-only constructs.**
Representative signals from a sequencing run containing only template DNA (i.e., without conjugated peptide or threading sequence). No reproducible low-amplitude intervals, characteristic of peptide-containing constructs, were detected.

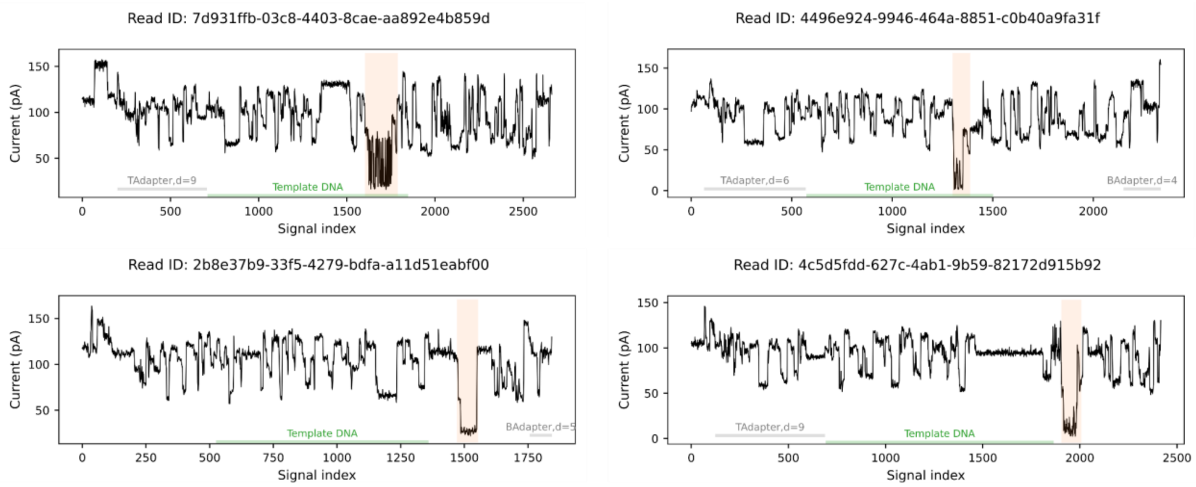


**Supplementary Figure 4 | False-positives from control experiment with template-only constructs.**
Representative signal traces from a sequencing run containing only template DNA (without conjugated peptide or threading sequence). A subset of traces is erroneously segmented as peptide-linked regions (PLRs); however, these exhibit distinct signal characteristics compared to true PLRs.

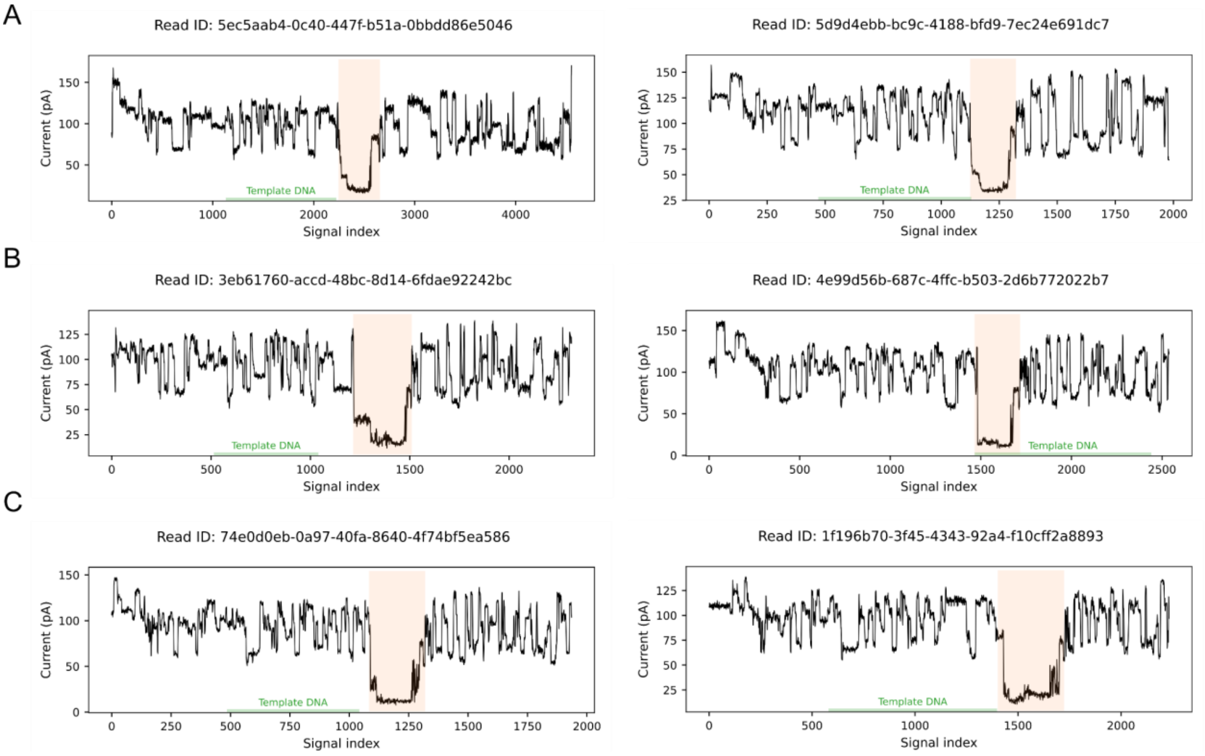


**Supplementary Figure 5 | PLR traces from βCAT-D (A), βCAT-L (B), and βCAT-W constructs (C).**

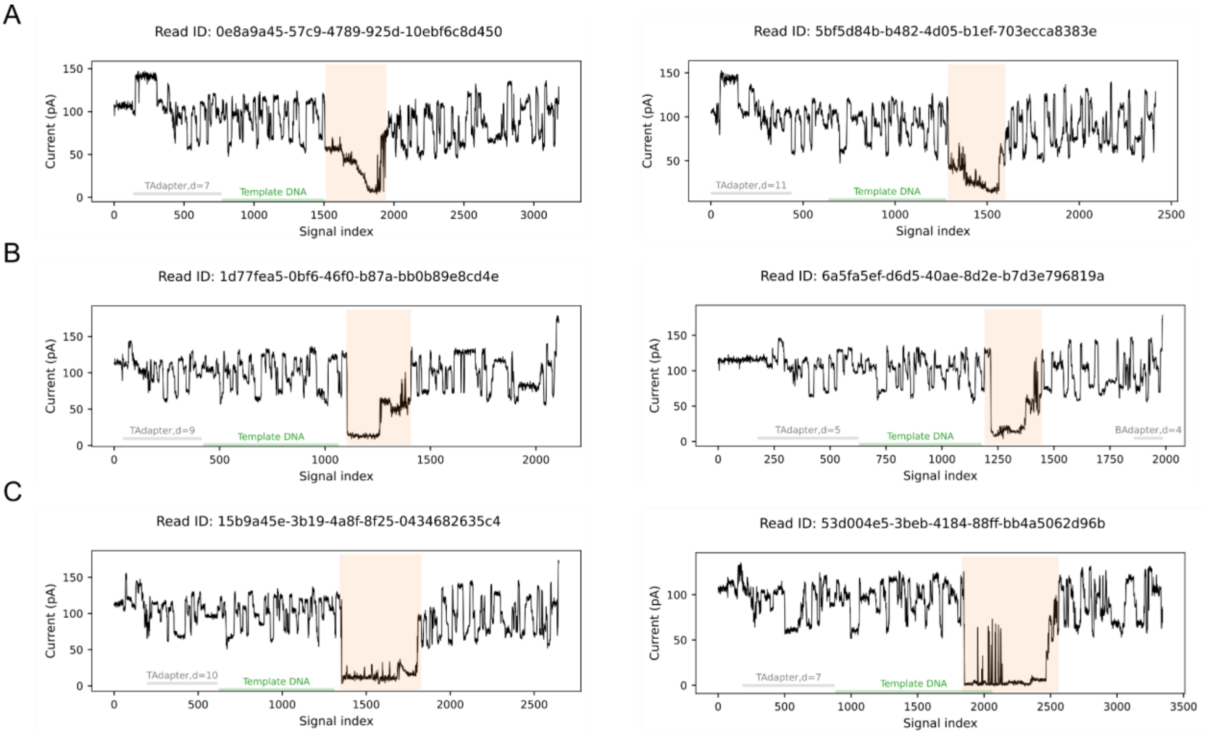


**Supplementary Figure 6 | PLR traces from BCAR3 (A), SZ-large (B), and ßCAT-30 (C).**

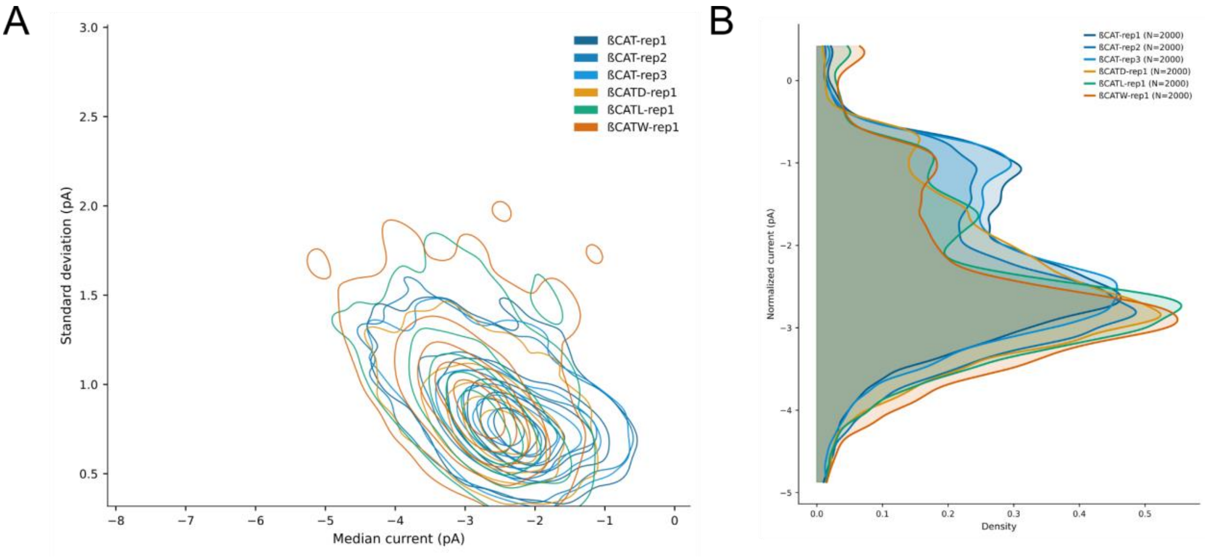


**Supplementary Figure 7 | Overlap of signal distributions across βCAT peptide variants.**
**(A)** Contour plots of signal distributions for four peptide variants, with median current (pA) on the x-axis and standard deviation (pA) on the y-axis, showing substantial overlap between variants. **(B)** Kernel density estimate (KDE) plots of ionic current distributions for βCAT, βCAT-D, βCAT-W, and βCAT-L. The x-axis represents ionic current (pA), and the y-axis indicates probability density.

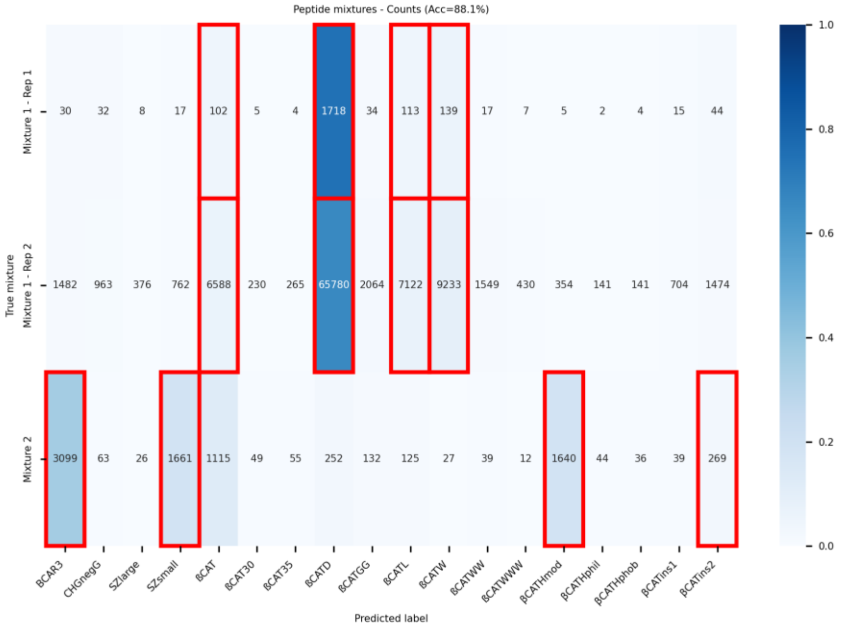


**Supplementary Figure 8 | Confusion matrix for peptide mixture experiments.**
Confusion matrix showing absolute prediction counts. Red rectangles indicate the ground truth peptides in each mixture.

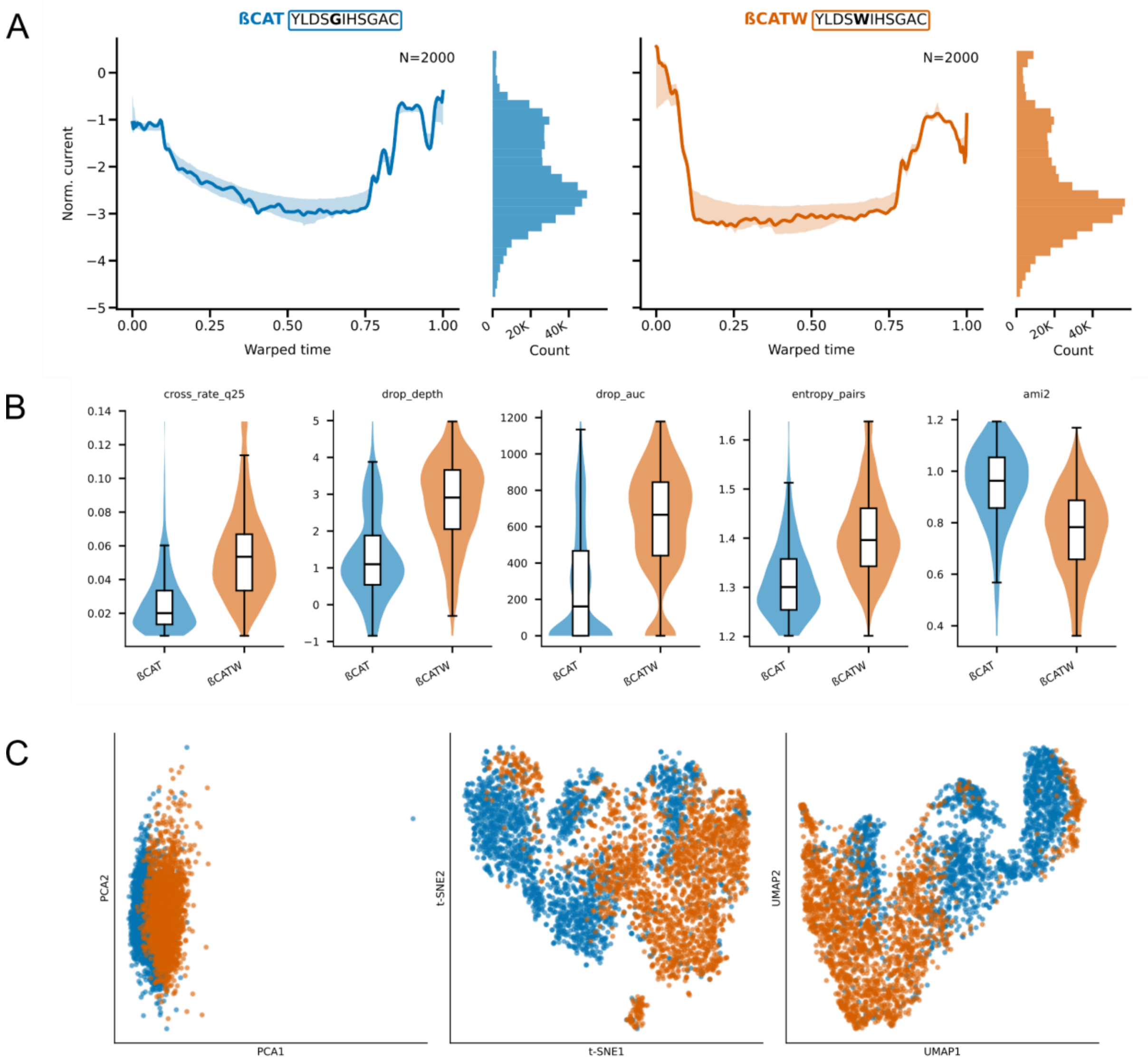


**Supplementary Figure 9 | Signal comparison between βCAT and βCAT-W.**
**(A)** Dynamic time warping (DTW) alignment of median signals for βCAT (blue) and βCAT-W (orange). Solid lines represent class median signals, with shaded regions indicating the standard deviation. The adjacent histogram shows the distribution of signal values across all measurements. **(B)** Box plots of five key signal features for βCAT and βCAT-W, highlighting systematic differences between variants. **(C)** Two-dimensional projections (PCA, t-SNE, UMAP) based on 30 extracted features for βCAT and βCAT-W.

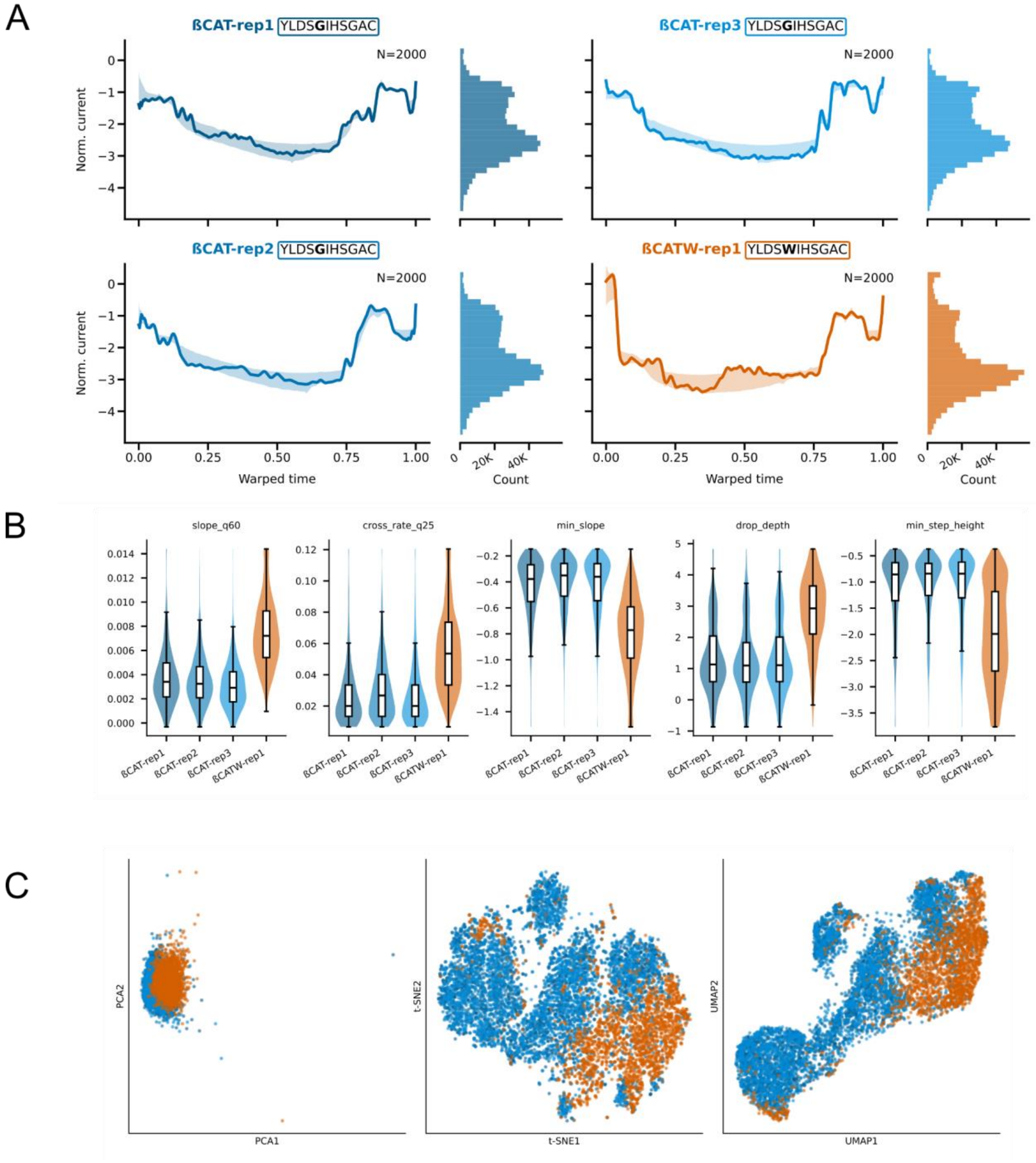


**Supplementary Figure 10 | Signal comparison between repeated measurements of βCAT and βCAT-W.**

**(A)** Dynamic time warping (DTW) alignment of median signals for βCAT-rep1, βCAT-rep2, βCAT-rep3, and βCAT-W. Solid lines represent class median signals, with shaded regions indicating the standard deviation. The adjacent histogram shows the distribution of signal values across all measurements. **(B)** Box plots of five key signal features across variants, highlighting systematic differences between βCAT and βCAT-W while illustrating consistency among repeated βCAT measurements. **(C)** Two-dimensional projections (PCA, t-SNE, UMAP) based on 30 extracted features for βCAT-rep1, βCAT-rep2, βCAT-rep3, and βCAT-W.

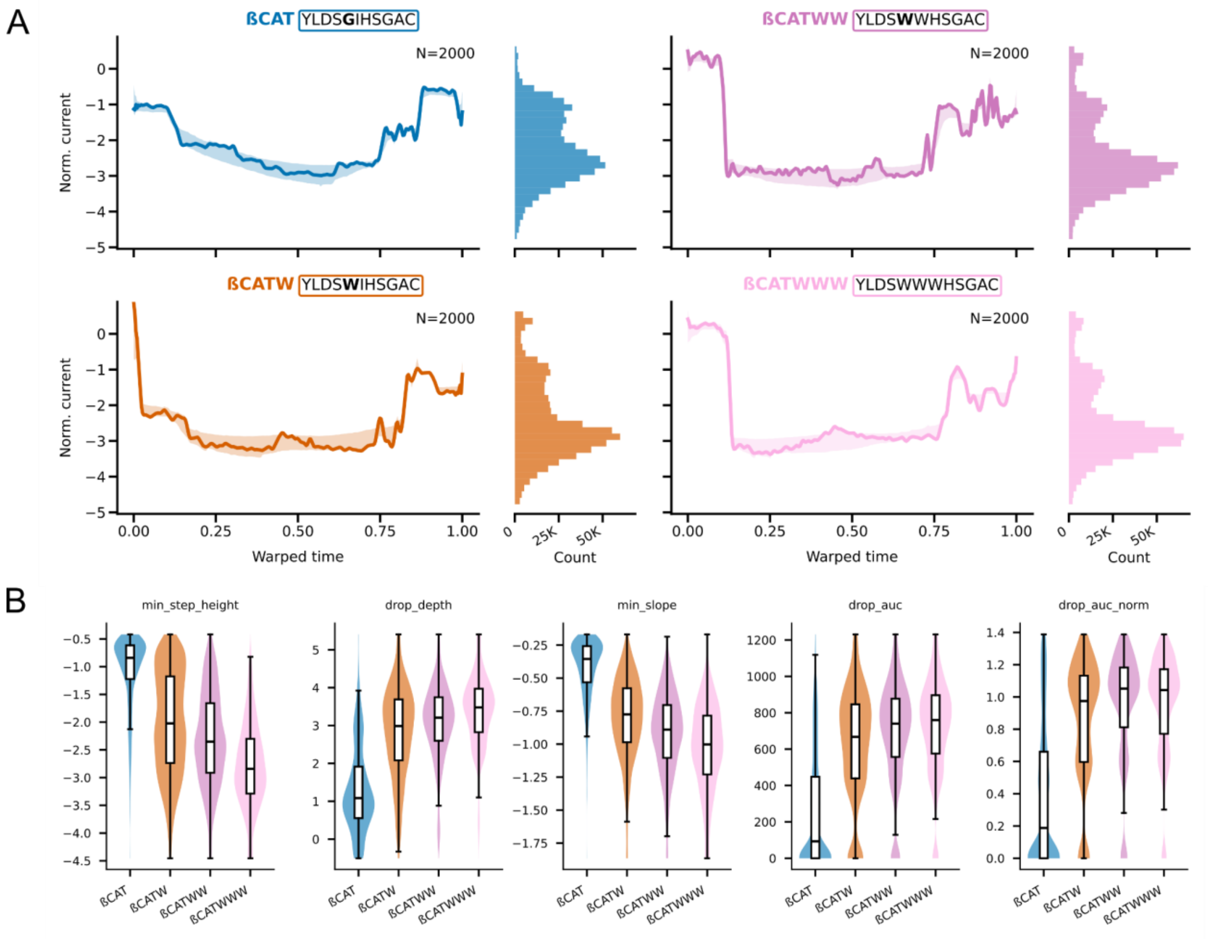


**Supplementary Figure 11 | Signal comparison between βCAT, βCAT-W, βCAT-WW, and βCAT-WWW**

**(A)** Dynamic time warping (DTW) alignment of median signals for βCAT, βCAT-W, βCAT-WW, and βCAT-WWW. Solid lines represent class median signals, with shaded regions indicating the standard deviation. The adjacent histogram shows the distribution of signal values across all measurements. **(B)** Box plots of five key signal features for each variant, highlighting systematic differences between classes.

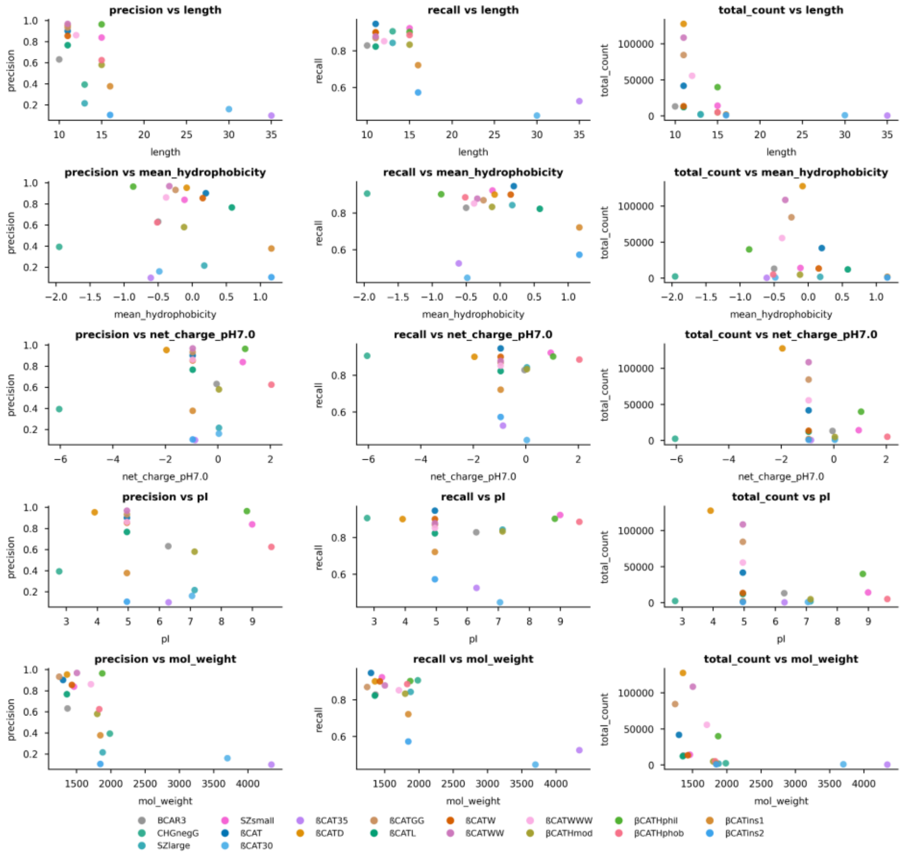


**Supplementary Figure 12 | Peptide physicochemical properties and corresponding classification performance metrics.**

Scatter plot grid showing peptide properties (length, mean Kyte-Doolittle hydrophobicity, net charge at pH 7, estimated isoelectric point, molecular weight; X) versus classification performance metrics (precision, recall, total PLRs per peptide; Y). Each marker represents a peptide from the validation set, with metrics aggregated as the median across replicate measurements, and is colored by peptide identity.

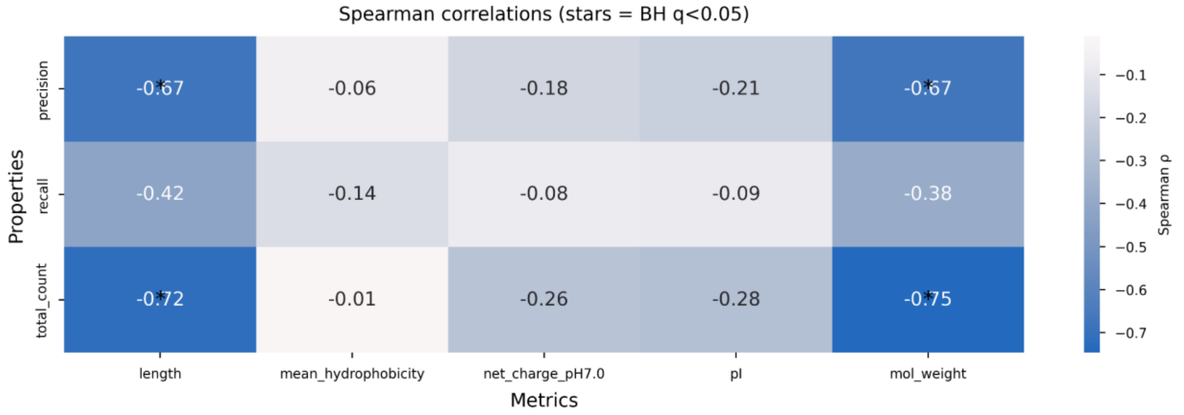


**Supplementary Figure 13 | Spearman correlations between peptide physicochemical properties and classification metrics.**

Heatmap summarizing Spearman rank correlations (ρ) between peptide properties (rows) and classification performance metrics (columns). Cells marked with an asterisk indicate statistical significance (q < 0.05) after Benjamini-Hochberg FDR correction .

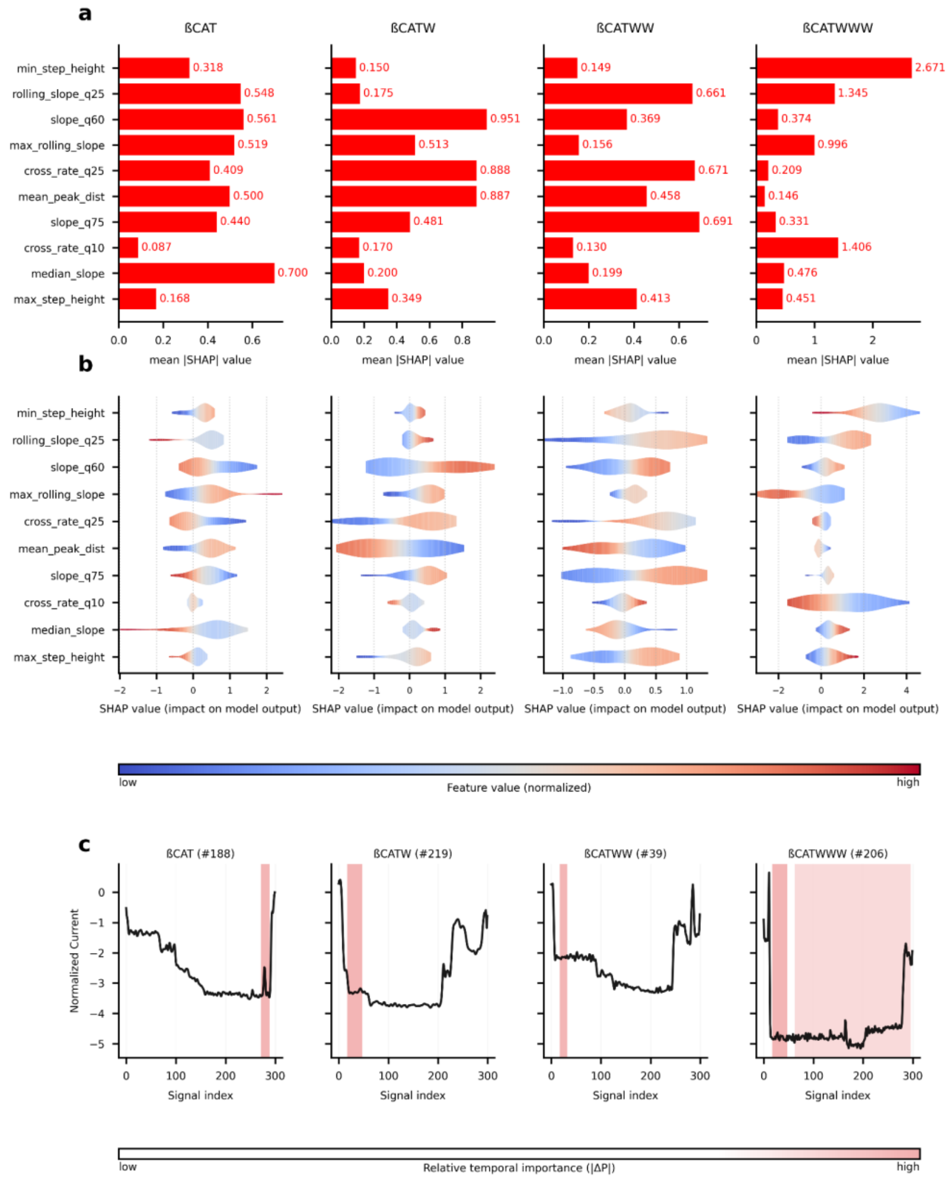


**Supplementary Figure 14 | Model interpretability and error analysis for W-variants of ßCAT.**

Each column corresponds to one peptide (ßCAT, ßCAT-W, ßCAT-WW and ßCAT-WWW), showing peptide-specific global, local and temporal feature importance**. (A)** Global SHAP feature importance showing the mean absolute SHAP value of the ten most influential features for each peptide, based on the global feature ranking across all classes. Higher values indicate stronger overall contribution to the model's predictions. **(B)** Local SHAP feature importance visualized as beeswarm plots for the same set of features. Each point represents a single prediction, where color denotes the corresponding normalized feature value (blue for low, red for high). Positive SHAP values indicate a feature increases the predicted probability for a given peptide, while negative values indicate the opposite effect. **(C)** Temporal feature importance for representative examples. Black traces show the raw nanopore current signals, and red-highlighted regions mark segments whose occlusion (replacement with current trace median) causes the largest change in predicted class.

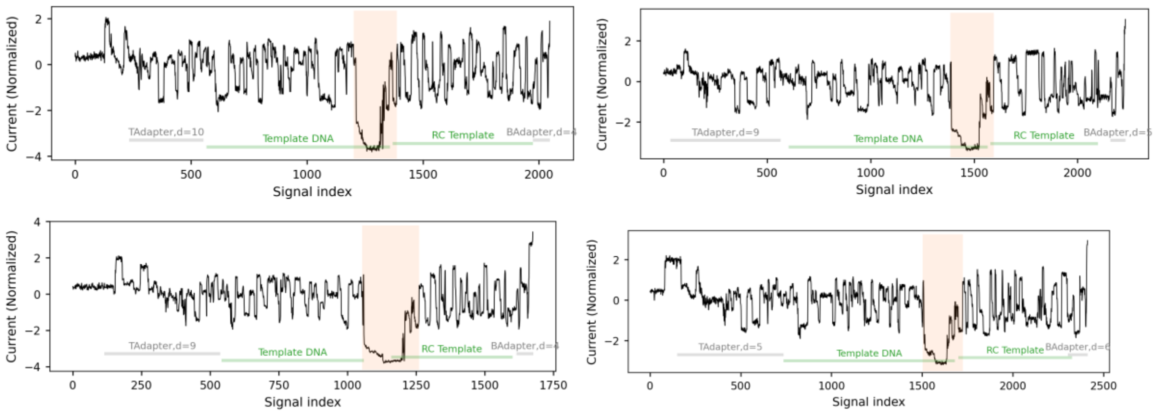


**Supplementary Figure 15 | Forward-reverse complement template chimeras identified in βCAT.**

Representative current traces showing alignment of the forward template segment followed by alignment to the reverse complement (RC) of the same template within a single read. Signal-to-reference alignment was performed using Remora. The transition from forward template to template-RC occurs within a continuous translocation event, consistent with the chimeric configuration described in Supplementary Note 01.

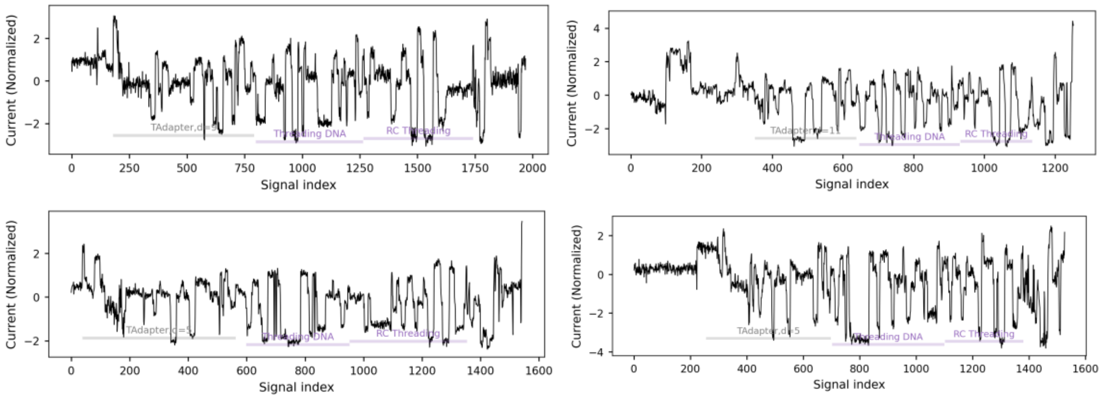


**Supplementary Figure 16 | Forward-reverse complement artefacts in reads aligning exclusively to threading DNA.**

Representative current traces from reads that align exclusively to the threading DNA segment, originating from sequencing runs in which template, threading, and peptide molecules were present in the mixture. Signal-to-reference alignment using Remora shows forward alignment of the threading segment followed by alignment to its reverse complement (RC) within the same read. No peptide-linked region (PLR) or current drop is observed, indicating that the forward-RC configuration occurs independently of peptide insertion.

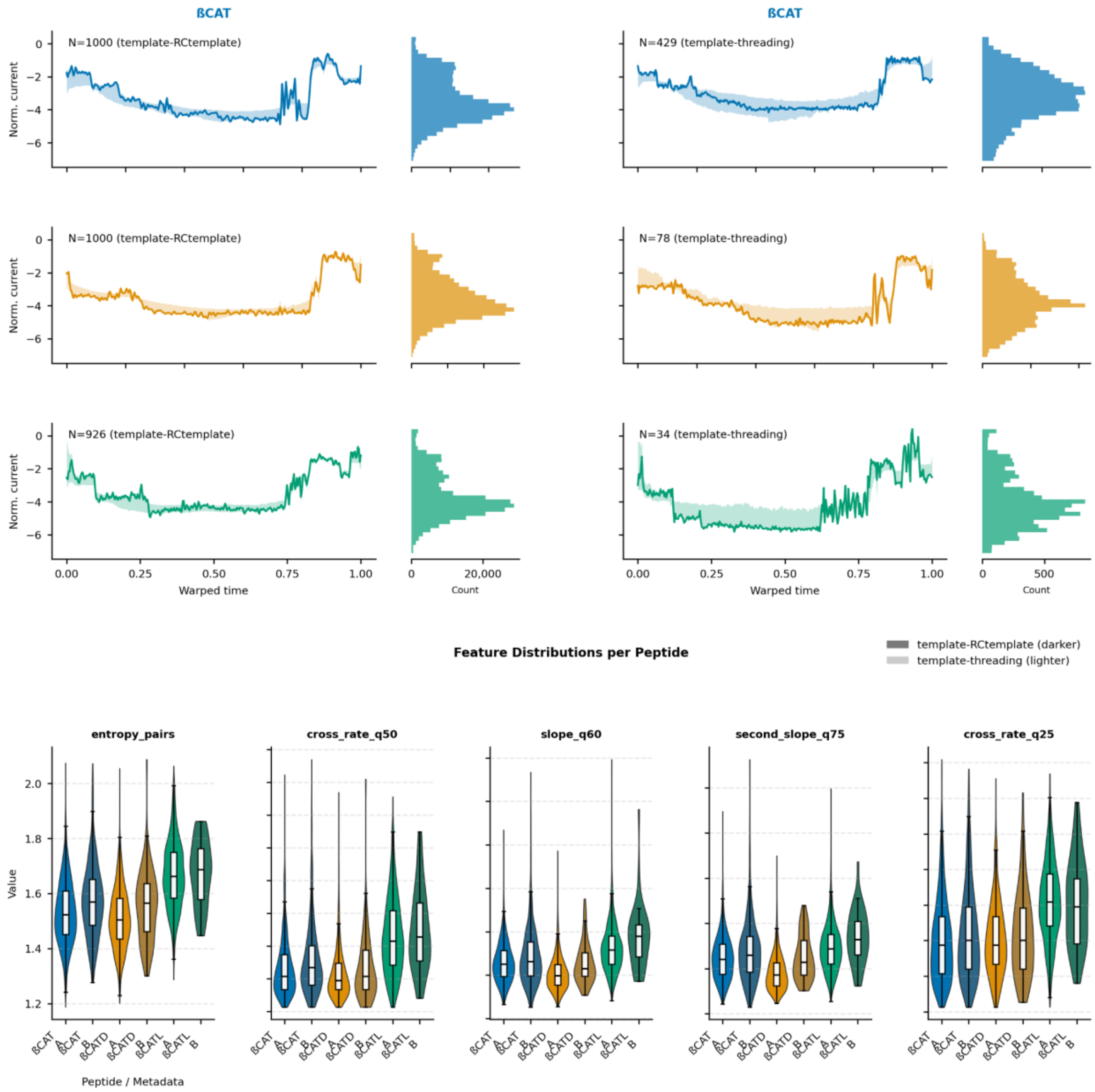


**Supplementary Figure 17 | Comparison of signal and features of template-RCtemplate and template-threading alignments.**

To evaluate whether PLR signals are comparable across alignment configurations, we compared PLRs fully aligned to template-RCtemplate and template-threading using identical preprocessing. **(Top)** Representative current signals for βCAT, ßCAT-D, and ßCAT-L are shown for template-RCtemplate (left) and template-threading (right). The dark solid trace is the DTW medoid signal for each class (the exemplar minimizing total DTW distance to all other class signals). Shaded bands indicate the interquartile range (25th-75th percentile) of signals after DTW alignment to the class medoid. Adjacent histograms summarize current-value distributions across all traces per class and alignment setting. Signal-display limits were set using the 1st-99th percentile range for visualization. **(Bottom)** Distributions of the five most discriminative signal features are shown for each peptide and alignment configuration. Violin plots with overlaid boxplots summarize central tendency and variability. Darker colors correspond to template-RC-template, and lighter colors to template-threading.

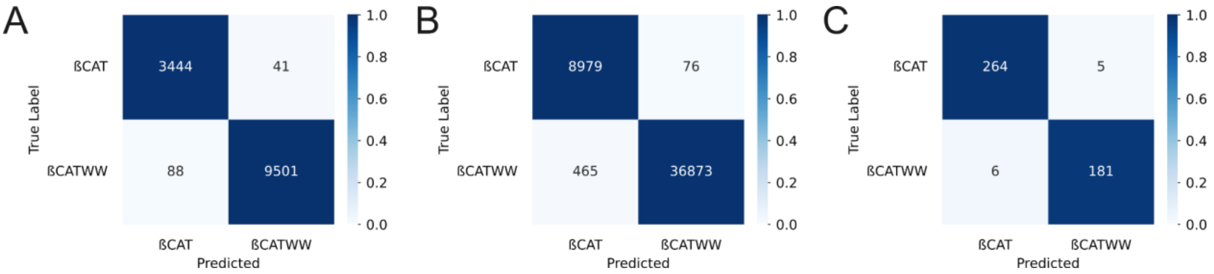


**Supplementary Figure 18 | Classification performance across alignment configurations for ßCAT and ßCAT-WW.**

Supervised classification with InceptionTime was trained exclusively on PLRs derived from reads fully aligning to the template-RC-template configuration. **(A)** Confusion matrix (absolute counts) for validation data consisting of template-RC-template PLRs. **(B)** Confusion matrix (absolute counts) for test data consisting of template-RC-template PLRs.**(C)** Confusion matrix (absolute counts) for test data consisting of template-threading PLRs.

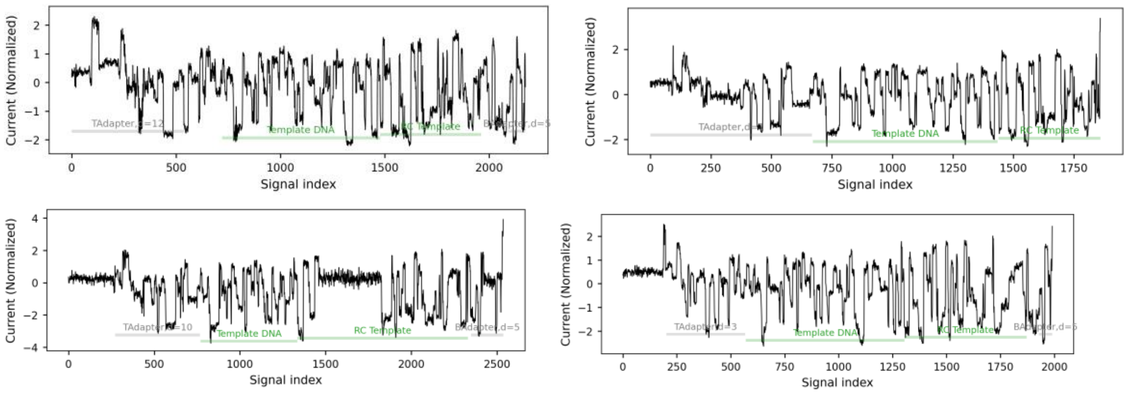


**Supplementary Figure 19 | Current traces from peptide-free control runs.**

Representative current traces from sequencing runs containing template DNA only (i.e., no threading strands or peptides added to the sequencing mixture). Experimental conditions and library preparation were otherwise identical to peptide-containing runs. Only canonical DNA translocation signals are observed, with no extended low-amplitude PLRs, supporting the peptide dependence of the PLR signature.

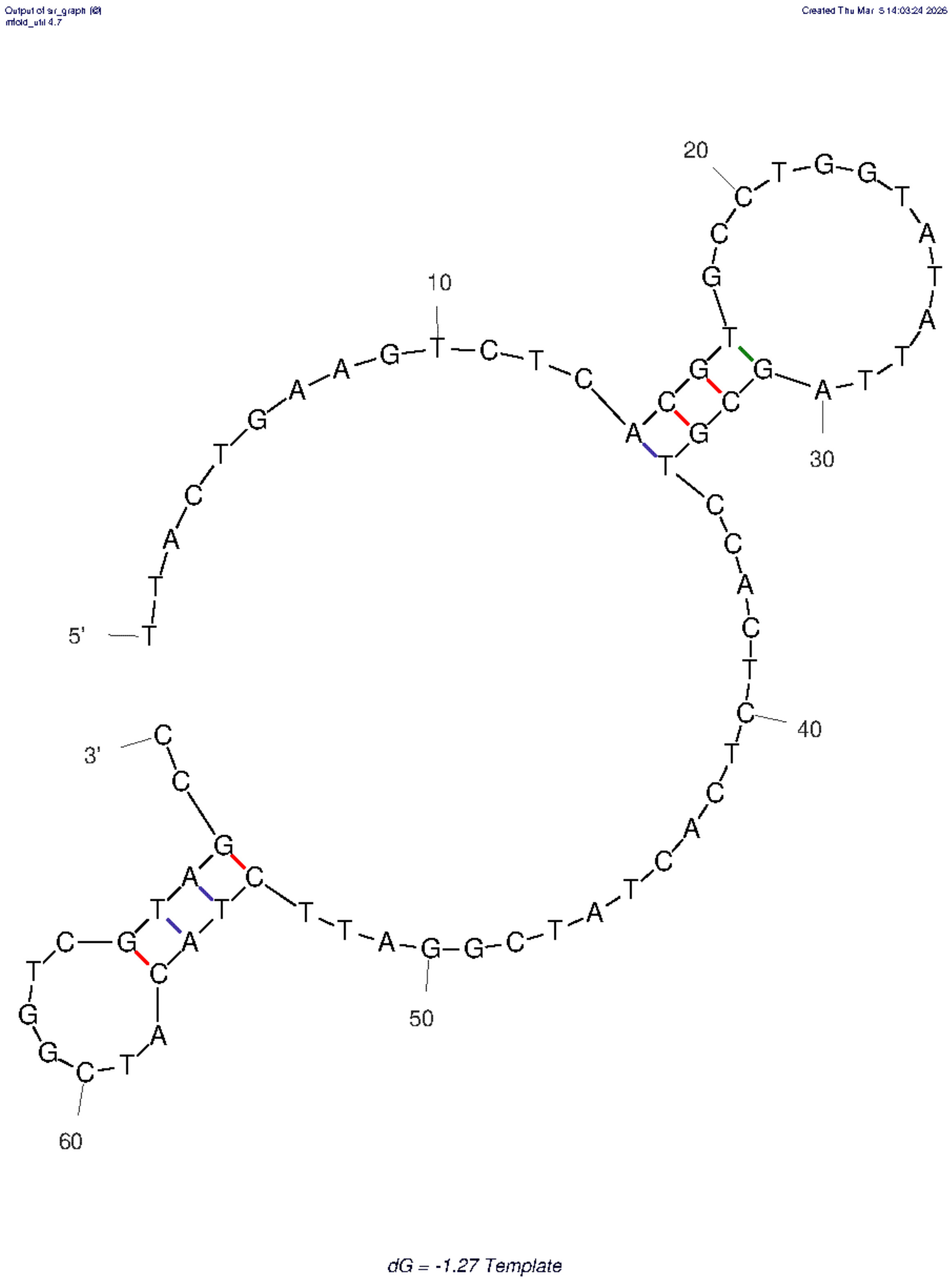


**Supplementary Figure 20 | Secondary structure prediction of the template DNA.** Predicted secondary structure of the single-stranded template DNA computed using UNAFold Mfold Webserver. Stable hairpin structures are observed near the position corresponding to the forward-reverse complement transition site identified in chimeric reads (Supplementary Note 01), supporting a potential foldback-based mechanism.

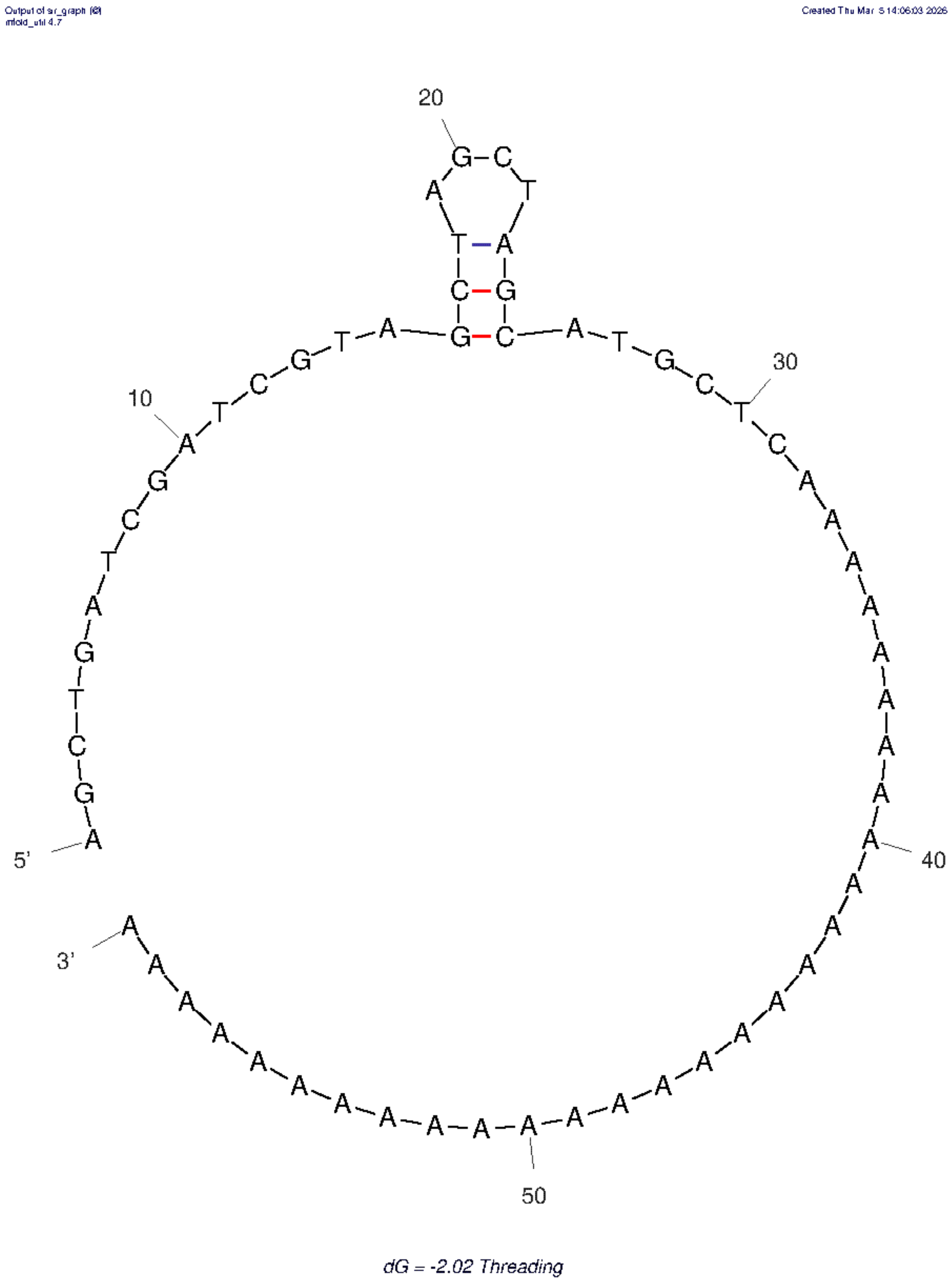


**Supplementary Figure 21 | Secondary structure prediction of the threading DNA.** Predicted secondary structure of the single-stranded threading DNA computed using UNAFold Mfold Webserver. Stable hairpin structures are observed near the position corresponding to the forward-reverse complement transition site identified in chimeric reads (Supplementary Note 01), supporting a potential foldback-based mechanism.

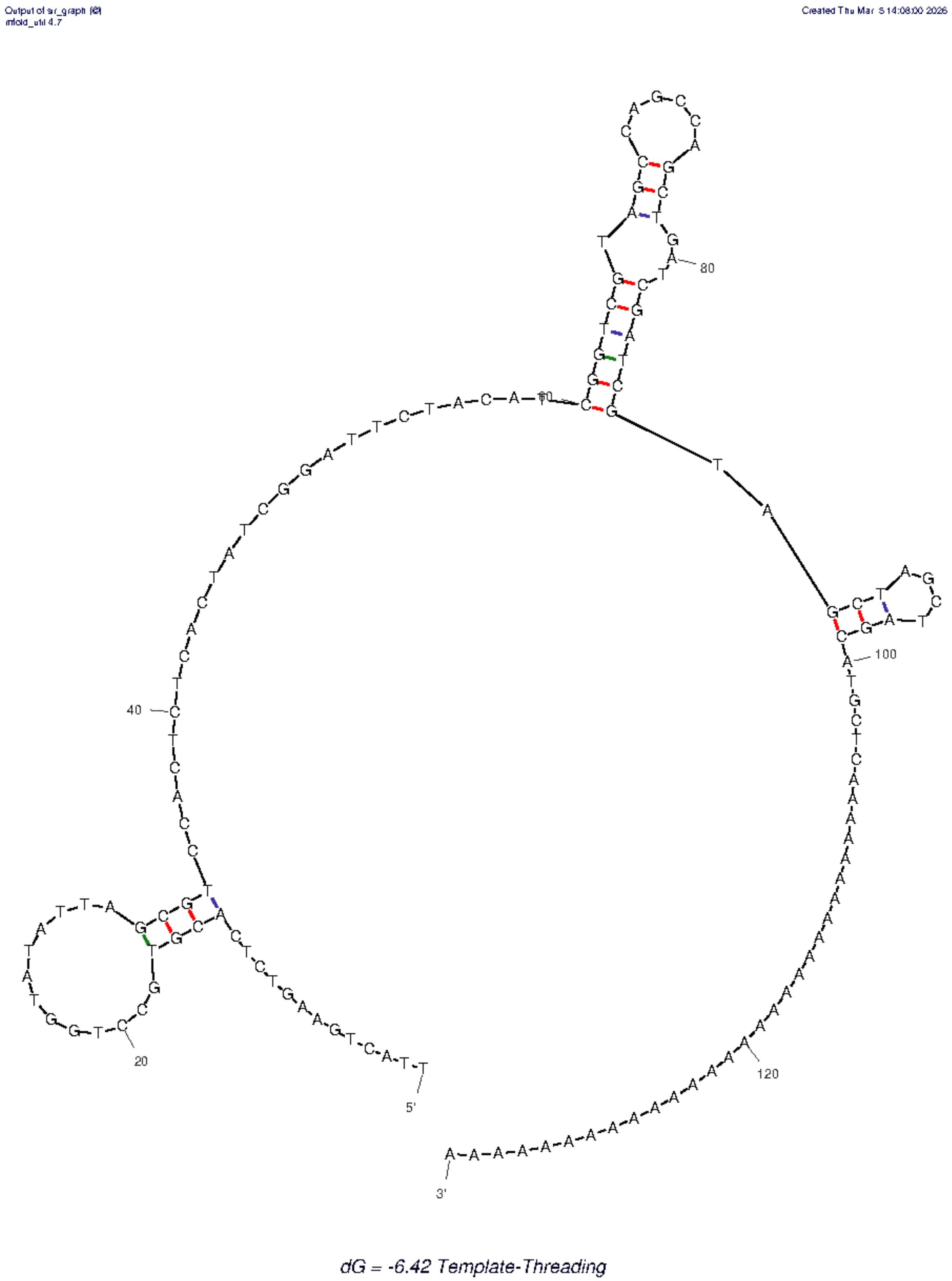


**Supplementary Figure 22 | Secondary structure prediction of the template-threading construct.**
Predicted secondary structure of the single-stranded template-threading DNA construct computed using UNAFold Mfold Webserver.
